# RNA-protein interactome at the Hepatitis E virus internal ribosome entry site

**DOI:** 10.1101/2022.04.11.487827

**Authors:** Shiv Kumar, Rohit Verma, Sandhini Saha, Ashish Agrahari, Shivangi Shukla, Anurag, Tushar Kanti Maiti, Shailendra Asthana, C.T. Ranjith-Kumar, Milan Surjit

## Abstract

Multiple processes exist in a cell to ensure continuous production of essential proteins either through cap-dependent or cap-independent translation processes. Viruses depend on the host translation machinery for viral protein synthesis. Therefore, viruses have evolved clever strategies to utilize the host translation machinery. Earlier studies have shown that genotype 1-Hepatitis E virus (g1-HEV) utilizes both cap-dependent and cap-independent translation machineries for its replication and proliferation. Cap-independent translation in g1-HEV is driven by an eighty seven nucleotide-long RNA element which acts as a noncanonical, internal ribosome entry site like (IRESl) element. Here, we have identified the RNA-protein interactome of the HEV IRESl element and characterized the functional significance of some of its components. Our study reveals indispensable roles of host ribosomal protein RPL5 and DHX9 (RNA helicase A) in mediating efficient translation from the IRESl element and establish the function of HEV IRESl as a bonafide internal ribosome entry site.

**Author summary:** Protein synthesis is a fundamental process for survival and proliferation of all living organisms. Majority of cellular proteins are produced through cap-dependent translation. Cells also utilize a variety of cap-independent translation processes to synthesize essential proteins during stress. Viruses depend on the host cell translation machinery to synthesize their own proteins. Hepatitis E virus is a major cause of hepatitis worldwide. The viral genome is a capped positive strand RNA. Viral non-structural and structural proteins are synthesized through a cap-dependent translation process. An earlier study from our laboratory reported the presence of a fourth ORF in genotype 1-HEV, which produced the ORF4 protein using a cap-independent internal ribosome entry site-like (IRESl) element. In the current study, we identified the host proteins that associate with the HEV-IRESl RNA and generated the RNA-protein interactome. Through a variety of experimental approaches, our data proves that HEV-IRESl is a bonafide internal ribosome entry site.

## Introduction

Protein synthesis is an essential process for survival and growth of all organisms. Viruses depend on the host cell machinery for synthesis of their proteins. Proteins are synthesized by the process of translation, which involves four distinct steps such as initiation, elongation, termination and recycling of the ribosomes. In eukaryotes, under normal conditions, majority of mRNA are translated by recognition of the m^7^G (5’)_PPP_(5’)N(cap) structure present at the 5’-end of the mRNA, by the process of cap-dependent translation. Many cellular and viral mRNAs are translated by the cap-independent mechanisms, by using distinct *cis*-acting regulatory elements, such as the cap-independent translational enhancers (CITEs), m^6^A modification of the mRNAs or the internal ribosome entry sites (IRES). Initiation of the cap-dependent translation occurs through a ribosomal scanning mechanism, in which the 48S initiation complex moves along the mRNA from the m^7^G cap in the 5’ to 3’ direction to locate a suitable AUG initiation codon. Following AUG recognition, through a series of steps, 80S ribosome complex is assembled on the mRNA with met-tRNAi bound to the AUG codon at the P-site, leading to translation elongation (1). The CITES are located in either 5’- or 3’-UTRs (untranslated regions) of the mRNAs and they initiate cap-independent translation by recruiting the initiation factors to the uncapped mRNAs, followed by 5’-end ribosomal scanning mediated identification of the initiation codon. Under conditions of stress, the cap-independent translation of many cellular mRNAs such as c-myc, Apaf-1, XIAP seem to occur through the above mechanism (2, 3). A variant of the eukaryotic initiation factor 4G (eIF4G) has been shown to be important for the cap-independent translation of the c-myc mRNA (4). Recently, yeast eIF4G1 was shown to mediate the cap-independent translation from the black beatle virus 5’-untranslated region (UTR) (5). Translation of 5’- or 3’-UTR m^6^A modified mRNAs also occur in a cap-independent manner (6,7,8).

Cap-independent, IRES-mediated translation has been reported in many positive-strand RNA viruses, which contain an uncapped genomic RNA. 5’-UTR in the genomic RNA of these viruses contain highly structured RNA elements, which directly recruit the initiation factors and promote translation through a scanning independent process, with the exception of the type I IRES, which depends on the ribosomal scanning process. IRES have been divided into 5 major types based on their mode of ribosome recruitment and RNA structure. The type I and type II IRES are found in the Picorna viruses such as the Polio virus (PV) and the Foot and Mouth Disease virus (FMDV), respectively. The PV IRES harbors six stem loops designated as domain I to VI. The Domain I forms unique clover leaf structure and is critical for replication of both the positive and the negative sense RNA. The domains II to VI are responsible for the PV IRES activity. During PV infections, viral 2A^pro^ cleaves the eIF4E binding N-terminal domain of the eIF4G without affecting its eIF3/eIF4A binding property. Stable association of the eIF4G with the PV IRES domain V promotes binding of other initiation factors, leading to formation of the 43S preinitiation complex. The FMDV IRES is a well-studied example of the type II IRES. The domain IV of the FMDV IRES binds with scaffold protein eIF4G. The 3C^pro^ and L^pro^ of FMDV cleave the eIF4G similar to the 2A^pro^ of the type I IRES. Notably, the FMDV IRES skips ribosomal scanning, instead, IRES proximal stem loop formation brings 84 nucleotides downstream AUG, close to the first AUG to start the translation by direct ribosome transfer. Other identified cis-acting elements for the type II IRES activity are the GNRA, RAAA and C-rich loop in the domain III (9). The type III IRES is present in the 5’-UTR of the Hepatitis A virus genome (10). It requires eIF4E binding for translation initiation (11). The type IV IRES have been reported in members of the Flaviviridae family, classified as the HCV (Hepatitis C virus)/ HCV-like IRES. The 5’-UTR of HCV contains four domains: the domain I and II are involved in the viral replication while the domains III and IV are involved in the IRES activity (12, 13). The HCV/HCV-like IRES are shorter in comparison to the type I and II IRES. The domains II and III contain several subdomains for interaction with the 40S ribosomal subunit. The type V IRES includes the long intergenic region (IGR) IRES, found between two open reading frames in the viral genomes and conserved in the dicistroviridae family (12, 13). The IGR IRES are the smallest IRES (∼180 nt long) elements consisting of three pseudoknots. It directly binds to the ribosomes and initiates translation with the alanine-tRNAi (ala-tRNAi) instead of the met-tRNAi, without involving the eIFs (14, 15).

Many viruses have evolved the ability to utilize both cap-dependent and cap-independent translation processes, depending on the state of the cell. Notably, although Dengue virus and Zika virus genomic RNA is usually translated through a cap-dependent mechanism, their 5’-UTRs also contain IRES-elements, which maintain viral translation under unfavorable conditions where cap-dependent translation is inhibited by the host (16,17,18). Similarly, genomic RNA of the Human Immunodeficiency Virus (HIV) type 1 and the Simian Immunodeficiency virus (SIV) contain IRES-elements in their 5’-leader sequences, which maintain viral translation when cap-dependent translation is inhibited (19,20,21,22). In addition, recent studies have identified functional IRES elements within open reading frames of viral protein coding genes. For example, five functional IRES-elements are located within the coding region of the non-structural and the structural proteins of the human rhinovirus 16, which are active under conditions of endoplasmic reticulum (ER) stress (23, 24). The Human cytomegalovirus pUL138 protein is translated by an IRES under conditions of stress (25). Further, coat protein of the Turnip crinkle virus (TCV) is translated through an unstructured IRES (26).

The Hepatitis E virus is a positive strand RNA virus of the family Hepeviridae. The viral genome is capped at the 5’-end followed by a short UTR of 25 nucleotides. The viral non-structural proteins are synthesized from the ORF1 located between 26-5107 nucleotides. Viral capsid protein is produced by the ORF2, located at the 3’-end between 5145-7127 nucleotides. A short 3’-UTR of 65 nucleotides is present downstream of the ORF2, followed by a poly-A tail (27). A third ORF is present between the ORF1 and ORF2, which produces ORF3 protein, an accessory protein that associates with the host Tumor susceptibility gene 101 and facilitates release of the progeny virus (28–30). The above genome organization is conserved among the 7 genotypes of the HEV. In contrast to zoonotic nature of some HEV genotypes, g1-HEV infection is restricted to the human. The g1-HEV does not replicate efficiently in mammalian cell lines (31).

Our earlier study demonstrated the coexistence of both cap-dependent and cap-independent translation in the g1-HEV (32). Although HEV non-structural and structural proteins are synthesized by a cap-dependent translation process, a protein essential for g1-HEV replication (ORF4 protein) is synthesized from an overlapping ORF located within the viral ORF1 through a cap-independent translation process. An 87-nucleotide conserved RNA regulatory element located upstream of the ORF4 coding region was found to drive cap-independent translation of the ORF4 protein. Interestingly, although ORF4 protein production in g1-HEV infected cells was markedly enhanced upon treatment of cells with ER stress-inducing compounds, HEV IRESl functioned efficiently irrespective of the ER stress inducer treatment in *in-vitro* translation assays or bicistronic reporter-based assays. Analysis of cap-independent translation driven by the HEV RNA regulatory element by bicistronic reporter assays along with site directed mutagenesis mediated mapping of the regulatory RNA sequence suggested the HEV RNA regulatory element to be an IRES-element. However, considering the small size of the regulatory element and its lack of strong homology with canonical IRES-elements, it was designated as an IRES-like (IRESl) element. The current study was designed to further explore the mechanism of HEV IRESl activity and verify its function as a bonafide internal translation initiation site. Host interaction partners of the HEV IRESl were identified and their importance in the IRESl-mediated translation was evaluated. Subsequent experiments showed association of the HEV IRESl RNA with actively translating ribosomes, supporting its role as a translation initiation element. Significance of these findings in validating the role of HEV IRESl as a bonafide internal ribosome entry site is discussed.

## Results

### Identification of host proteins that interact with the HEV IRESl element

Two independent experimental approaches (RaPID assay and Yeast three hybrid assay) were followed to identify the host proteins that interact with the HEV IRESl RNA. RNA-protein interaction detection assay (RaPID assay) involves biotinylation of IRES interacting proteins, followed by their identification by liquid chromatography with tandem mass spectroscopy (LC-MS-MS) (33). RaPID assay was optimized in our laboratory (34). 87 nucleotides of the g1-HEV genome (Genbank ID: AF444002.1) corresponding to the IRESl region was cloned into the pRMB vector between the BirA ligase binding stem-loop (RMB-SLI, RMB-SLII) sequences (Fig. 1A, 1C). Analysis of the secondary structure of the fusion RNA sequence using “Mfold” indicated that the HEV IRESl and the BirA ligase binding stem-loops retained their distinct folding characteristics (Fig. 1A). An unrelated sequence in the g1-HEV genome (618-738 nucleotides from the 5’-end, denoted as control RNA) was also cloned into the pRMB vector to be used as a control to monitor specificity of the assay. Biotin-untreated cells were used as an additional control and experiments were performed two times and samples of each experiment were run in triplicate.

**Figure 1.**
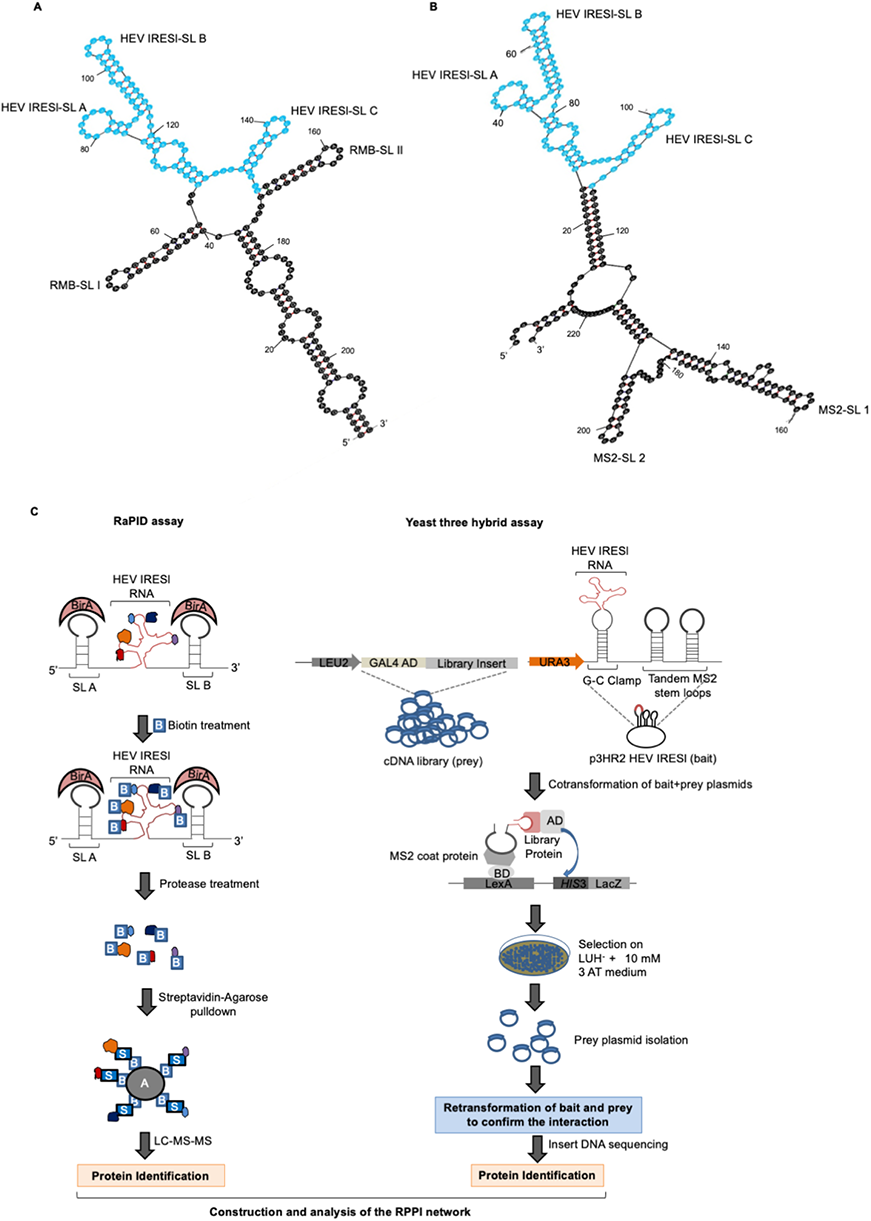
Identification of host interaction partners of the HEV IRESl element by RaPID and Yeast three hybrid assay. (A) Schematic of predicted secondary structure of the HEV IRESl RNA (highlighted in blue) fused to the BirA-binding RNA motifs (RMB-SL1 and RMB-SL2). Stem-loops of the HEV IRESl are denoted as SL A, SL B, SL C (B) Schematic of predicted secondary structure of the HEV IRESl RNA (highlighted in blue) fused to the MS2 coat protein-binding RNA motifs (MS2-SL1 and MS2**-**SL2). Stem-loops of the HEV IRESl are denoted as SL A, SL B, SL C (C) Work-flow to identify the HEV IRESl-binding host proteins. B: Biotin, A: Agarose, S: Streptavidin, AD: GAL4-Activation domain, BD: GAL4-DNA Binding domain, L: Leucine, U: Uracil, H: Histidine, 3-AT: 3-Amino 1, 2, 4 Triazole, “−”: Deficiency in the medium, “+”: Supplemented in the medium.

Quality of the MS data between replicates of different samples was checked by Pearson correlation analysis, which showed good correlation (average range: 0.2-1.0). Proteins with at least one biotinylated peptide with PEP (posterior error probability) score ≥ the median value in the Gaussian smoothing curve of each sample were selected for further analysis. Specific interaction partners of the HEV IRESl RNA were selected by subtracting the biotinylated HEV IRESl data set from the biotinylated HEV (618–738), unbiotinylated HEV (618–738) and unbiotinylated HEV IRESl datasets (Fig. 2A, Fig. S1, Table S1). The HEV IRESl RNA-binding protein dataset thus obtained was further analyzed to select only those proteins that were represented by one or more unique peptides and a “prot score” of 30 or more. Thus, a total of 43 unique proteins were identified as interaction partners of the HEV IRESl RNA (Fig. 2B, Table 1).

**Figure 2.**
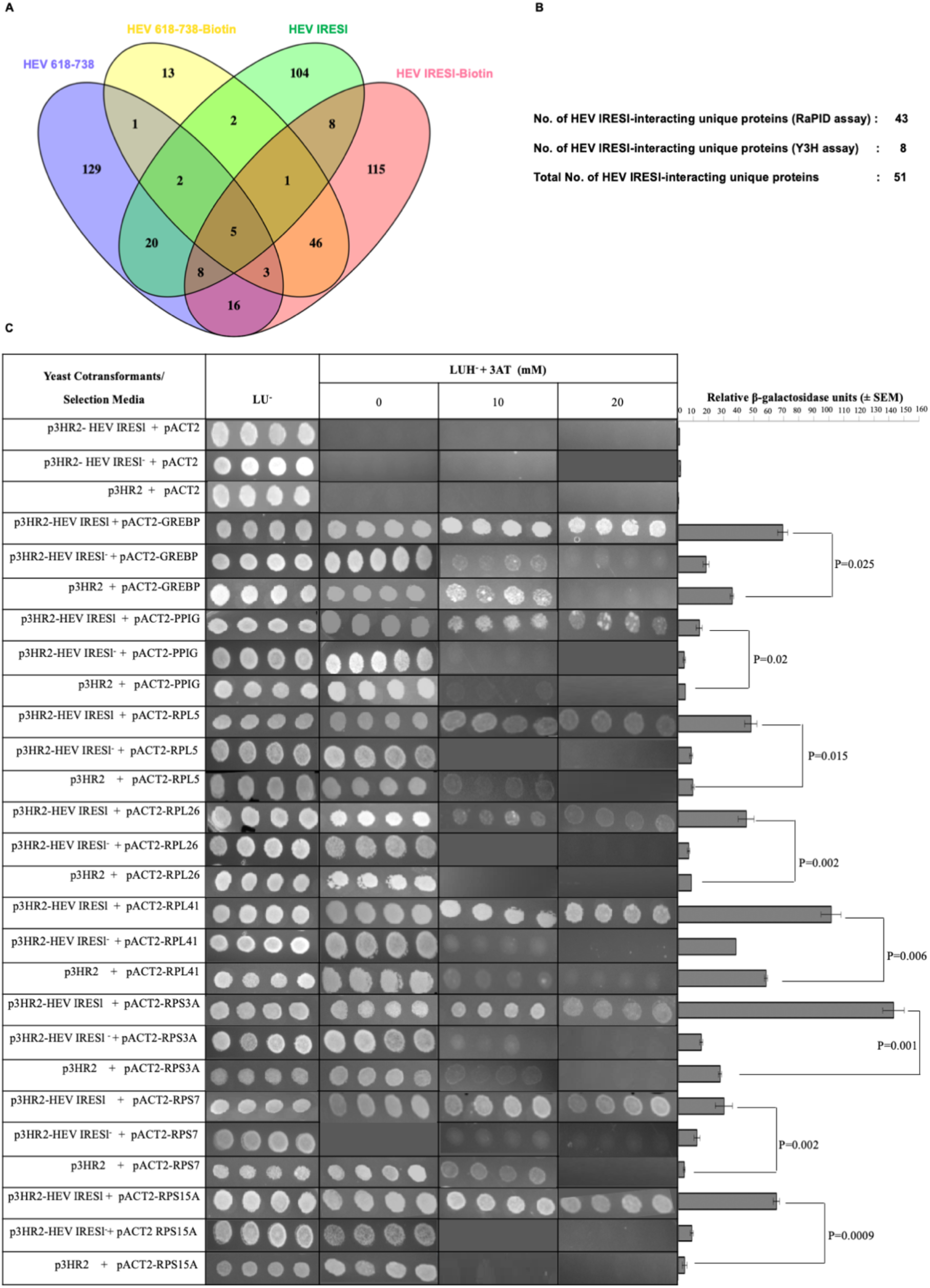
Identification of HEV IRESl RNA-binding proteins by RaPID and Y3H assay. (A) Venny analysis of the HEV IRESl RNA-binding proteins identified by the RaPID assay. (B) Summary of the HEV IRESl RNA-binding proteins identified by the RaPID and the Y3H assay. (C) Y3H assay mediated confirmation of the interaction between HEV IRESl RNA and the host proteins identified by screening of the human liver cDNA library. YBZ1 strain was transformed in the indicated combinations and plated on media lacking Leucine and Uracil (LU^-^). Four random colonies from each cotransformant plate were replica plated onto media lacking Leucine, Uracil, Histidine (LUH^-^) and supplemented with 5mM or 10mM 3-amino 1, 2, 4 Triazole (3-AT). Same colonies were used in liquid β-galactosidase assay. Relative β-galactosidase units are plotted as mean (± SEM).

**Table 1.**
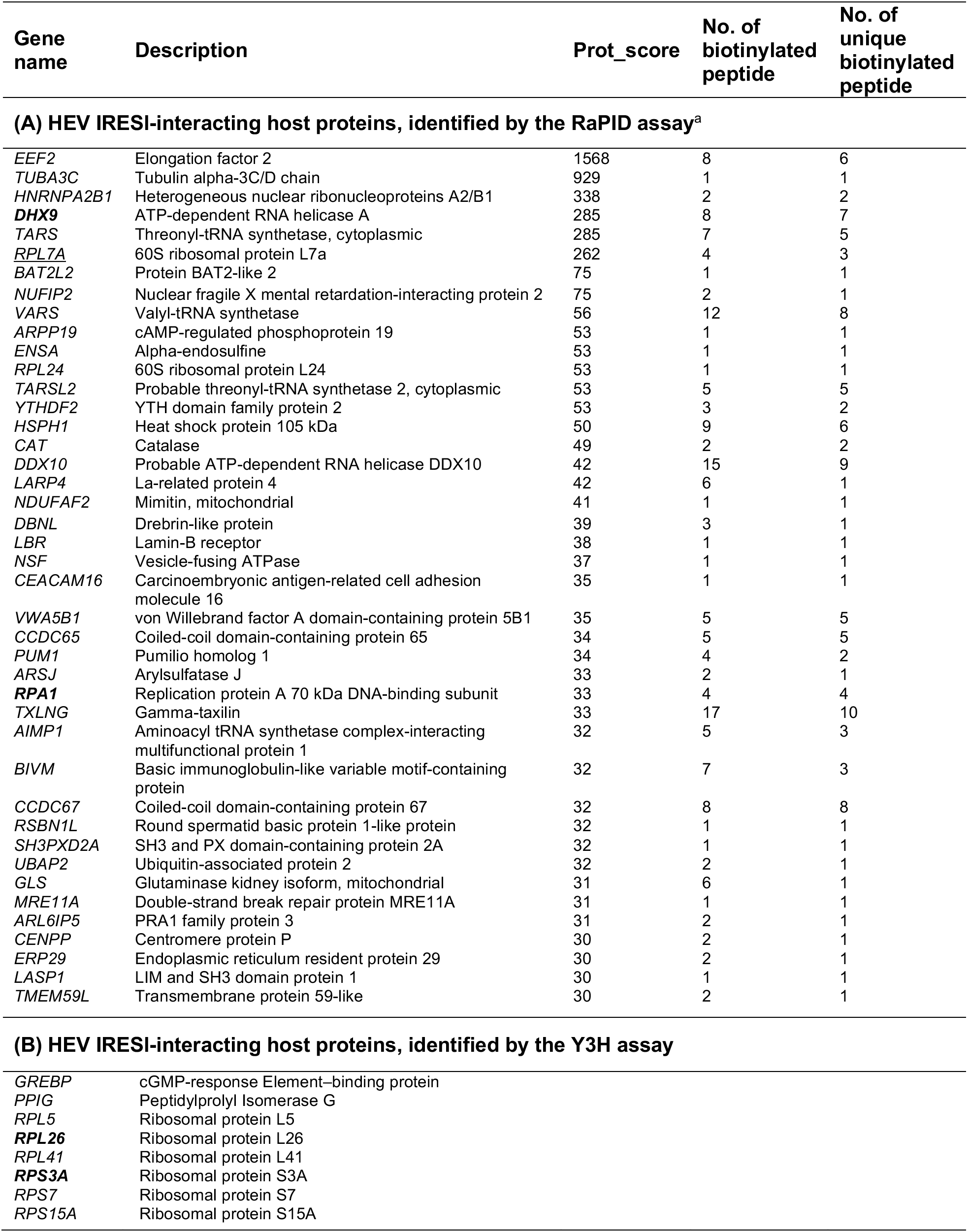
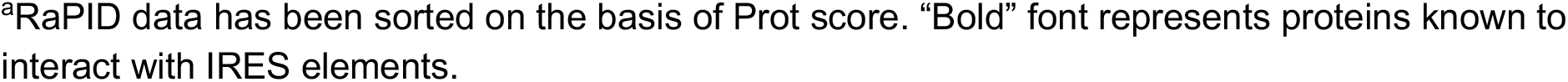
Host interaction partners of the HEV IRESl element

Since RaPID assay may have the limitation of not identifying all proteins buried inside the RNA-protein complex, yeast three hybrid (Y3H) assay was used as an alternate approach to unbiasedly identify the direct interaction partners of the HEV IRESl RNA. A human liver cDNA library was screened using the Y3H assay (Fig. 1C) (35, 36). In Y3H assay, the bait RNA is flanked by two copies of the MS2-coat protein-binding RNA element. Analysis of the RNA sequence containing fusion of the HEV-IRESl RNA and the MS2-coat protein binding stem loop RNA sequences using “Mfold” indicated that their secondary structure profiles remain unaltered in the fusion RNA (Fig. 1B). Self-activation of the *lacZ* and *HIS3* reporter genes by the HEV IRESl RNA was checked by cotransforming the plasmids encoding the HEV IRESl (p3HR2-HEV IRESl) and the GAL4 activation domain (pACT2) into chemically competent YBZ1 yeast cells, followed by selection of the cotransformants on the LUH^-^ medium and assay of the β-galactosidase activity. No growth of colonies was seen on the LUH^-^ plates neither β-galactosidase activity was observed, indicating that there was no self-activation of the reporter genes by the bait RNA (Fig. 2C). Next, the bait RNA expressing YBZ1 cells were transformed with the human liver cDNA library. A total of 4×10^6^ clones were screened and cotransformants were selected on the LUH^-^ medium supplemented with 10mM 3-amino 1,2,4 triazole(3-AT). Note that 3-AT is a competitive inhibitor of the Histidine biosynthesis pathway. Therefore, addition of 3-AT allows growth of the cotransformants that show strong interaction between the bait RNA and the prey protein. 395 colonies were obtained in the primary screening, out of which human liver cDNA insert could be detected in 285 colonies (Table S2). Restriction pattern analysis of the prey plasmid DNA isolated from the 285 colonies showed that unique cDNA insert was present in 75 plasmids. Analysis of the cDNA insert sequences identified 8 protein coding genes, which were in frame with the GAL4 activation domain (Table S2). Retransformation of those 8 plasmids along with the p3HR2-HEV IRESl plasmid showed that all of them were able to interact with the HEV IRESl (Fig. 2C). The eight proteins include 6 ribosomal proteins (RPL5, RPL26, RPL41, RPS3A and RPS7, RPS15A), PPIG (peptidylprolyl isomerase G) and GREBP (cyclic-GMP response element binding protein) (Table 1).

Further, specificity of the interaction of eight Y3H-identified proteins with the HEV IRESl was evaluated in two ways: (a) by measuring the interaction of those proteins with an RNA sequence complementary to the HEV IRESl (HEV IRESl^-^) (b) by measuring the interaction of those proteins with the known IRES sequences from the genotype 3A Hepatitis C virus (HCV) and the foot and mouth disease virus (FMDV). All sequences were cloned into the p3HR2 vector, upstream of the MS2 coat protein-binding stem-loop RNA sequences (Fig. S2). No significant interaction was observed between HEV IRESl^-^ RNA and the host proteins (Fig. 2C). Y3H analysis revealed that only RPS3A interacts with equal strength with all the three RNA sequences (HEV IRESl, FMDV IRES, HCV IRES) (Table S3 and Fig. 3). RPL41 and RPS15A interacts exclusively with the HEV IRESl RNA. While, RPL5 and RPS7 were found to interact strongly with the HEV IRESl RNA and weakly with the HCV IRES. RPL26 interacts moderately and strongly with the HEV IRESl and the HCV IRES, respectively, whereas it did not interact with the FMDV IRES. GREBP interacts strongly with the HEV IRESl and weakly with both FMDV IRES and HEV IRESl. PPIG also showed a difference in the strength of interaction among the three RNA sequences in the following order-HCV IRES: strong interaction, HEV IRESl: weak interaction and FMDV IRES: no interaction (Table S3 and Fig. 3).

**Figure 3.**
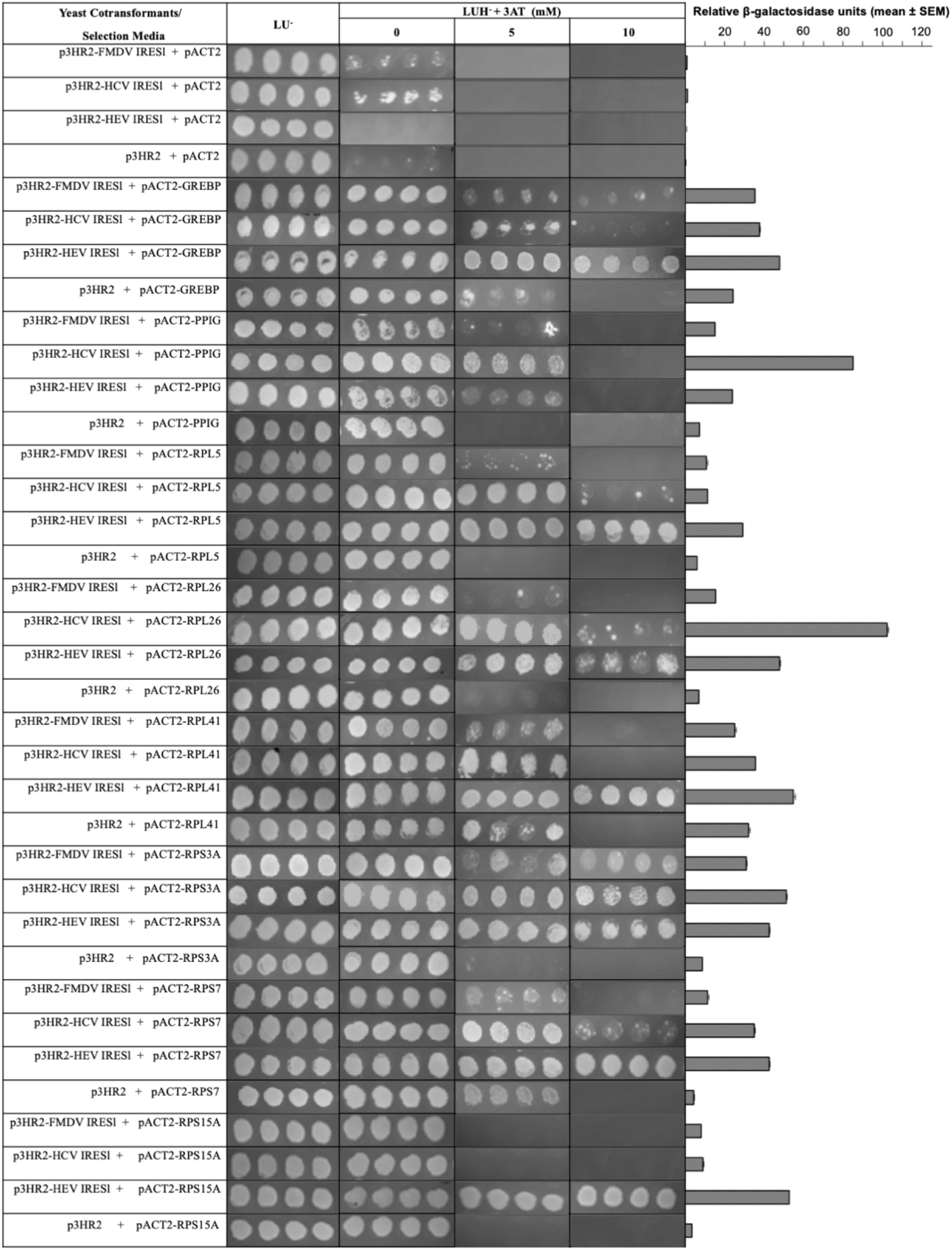
Assessment of the ability of the HEV IRESl-binding host proteins (identified by the human liver cDNA library screening) to interact with the HCV IRES and the FMDV IRES. YBZ1 strain was transformed in the indicated combinations and plated on media lacking Leucine and Uracil (LU^-^). Four random colonies from each cotransformant plate were replica plated onto media lacking Leucine, Uracil, Histidine (LUH^-^) and supplemented with 5mM or 10mM 3-amino 1, 2, 4 Triazole (3-AT). Same colonies were used in liquid β-galactosidase assay. Relative β-galactosidase units are plotted as mean (± SEM).

In order to further ascertain that HEV IRESl interaction partners identified by RaPID and Y3H are bonafide candidates, some of the interactions were further validated by an *in vitro* pull-down assay using *in vitro* transcribed biotinylated HEV IRESl RNA as bait. Non-biotinylated HEV IRESl and biotinylated control RNA (618-738 nts in g1-HEV genome) as well as only cell lysate (Mock) was used as controls to ensure specificity of the assay (Fig. 4A, 4B). All 10 proteins tested by the pull-down assay showed interaction with the HEV IRESl RNA (Fig. 4C). Six of these interactions were originally identified by Y3H assay (RPS3a, RPS7, RPL26, RPL41, PPIG, GREBP) and four were identified by RaPID assay (RPL5, RPL24, LARP4, DHX9) (Fig. 4C). These data further support that the proteins identified by RaPID and Y3H assay are bonafide interaction partners of the HEV IRESl RNA.

**Figure 4.**
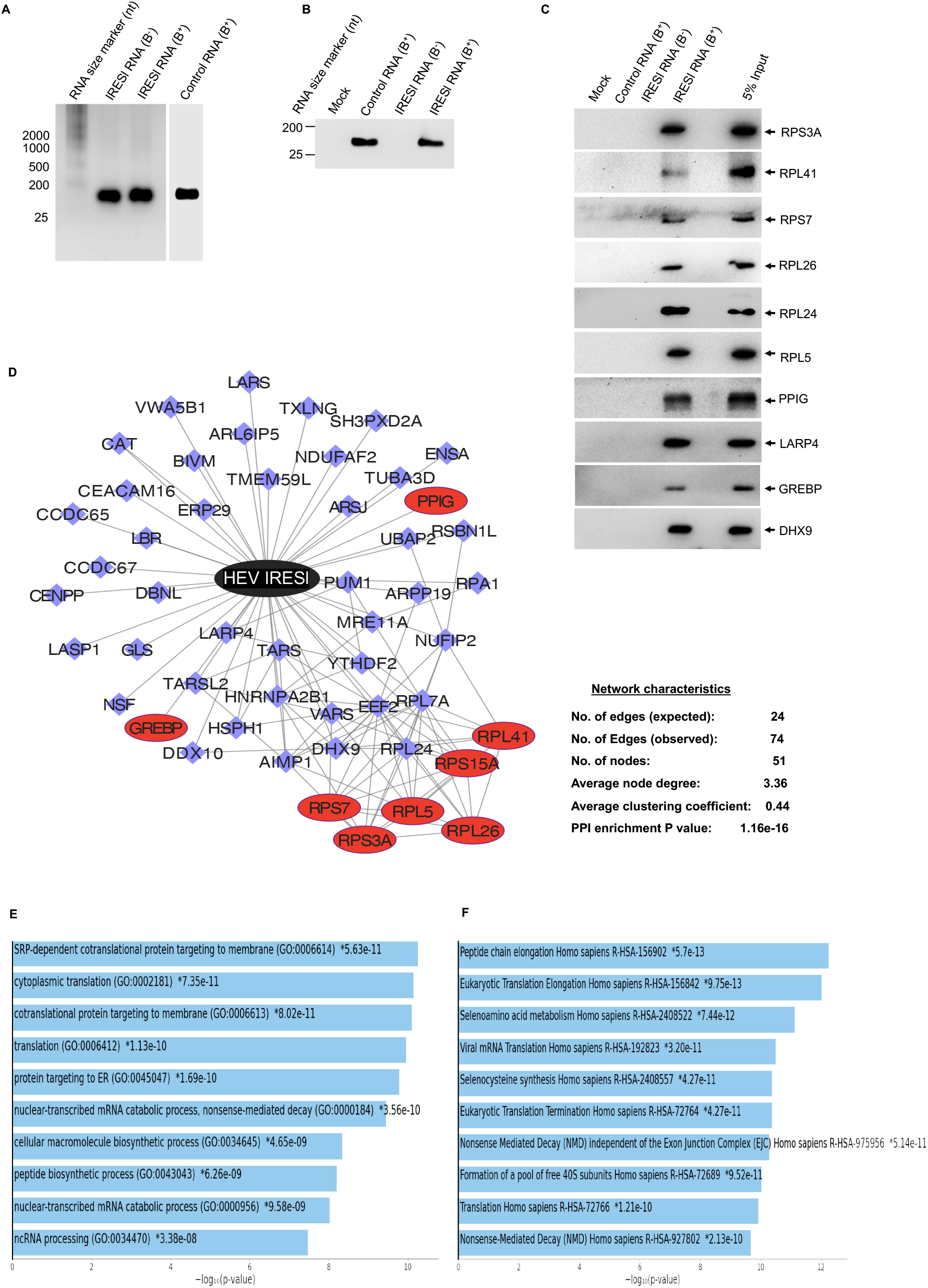
Confirmation of HEV IRESl RNA-protein interactions by biotinylated-RNA pull down assay and construction of the RNA-protein interaction network. (A) Formaldehyde-agarose gel electrophoresis and ethidium bromide staining mediated visualization of *in vitro* transcribed and purified non-biotinylated IRSEl RNA [IRESl RNA (B^-^)], biotinylated IRSEl RNA [IRESl RNA (B^+^)] and biotinylated control RNA [control RNA (B^+^)], as indicated. RNA size marker was resolved in parallel (lane 1). (B) Formaldehyde-agarose gel electrophoresis and ethidium bromide staining mediated visualization of eluates from streptavidin-pull down of samples containing the control RNA (B^+^), IRESl RNA (B^-^) and IRESl RNA (B^+^), as indicated. (C) Western blot analysis of the indicated proteins using aliquots of eluates from biotinylated-RNA pull down of samples containing the control RNA (B^+^), IRESl RNA (B^-^) and IRESl RNA (B^+^), as indicated. 5% input denotes 5% of the cell lysate used for incubation with the RNAs during the pull-down assay. (D) Schematic of the HEV IRESl RNA-protein interaction network. Black node denotes HEV IRESl RNA, blue and red nodes denote host proteins identified by the RaPID and the Y3H assay, respectively. Edges are represented by blue lines (E) Graphical representation of the top 10 Biological processes (sorted by *P*-values) enriched in the HEV IRESl RNA-protein interactome (F) Graphical representation of the top 10 Reactome pathways (sorted by *P*-values) enriched in the HEV IRESl RNA-protein interactome.

### Construction of the HEV IRESl RNA-host protein interaction network and analysis of the interactome

43 and 8 host proteins identified as interaction partners of the HEV-IRESl RNA by RaPID and Y3H assay, respectively, were clubbed and imported to Cytoscape to generate the RNA-Protein-Protein interaction (RPPI) network, as described earlier (29). Analysis of the different network characteristics revealed that the HEV IRESl-RPPI network contained 51 nodes and 74 edges, the average node degree being 3.36. Average clustering coefficient was 0.44 and the PPI enrichment P-value was 1.11e^-16^ (Fig. 4D). Above network parameters suggest that the HEV IRESl-RPPI network is highly connected and suitable for further downstream studies.

Next, Gene ontology (GO) and Reactome pathway analysis of the HEV IRESl-RPPI network was performed using the “Enrichr” tool to determine the significantly enriched processes/pathways. Proteins involved in translation and mRNA catabolic processes were enriched in the GO biological processes category, supporting involvement of the HEV IRESl element in the protein synthesis process (Fig. 4E, top 10 biological processes are shown). Similarly, top Reactome pathways include translation, eukaryotic translation elongation and seleno animo acid metabolism (Fig. 4F, top 10 Reactome pathways are shown). Analysis of the dataset using the gene set enrichment analysis (GSEA) tool also showed translation and peptide metabolic process as top hits, in agreement with the “Enrichr” output (Fig. S3A, S3B). Components of both small and large subunits of the ribosome such as RPS7, RPS3A, RPS15A and RPL5, RPL24, RPL26, RPL41 interact with the HEV IRESl. Components of the tRNA synthetase complex such as Threonyl-tRNA synthetase (TARS), Valyl-tRNA synthetase (VARS), Aminoacyl tRNA synthetase complex-interacting multifunctional protein 1 (AIMP1) and a component of the translation elongation complex (EEF2, eukaryotic elongation factor 2) also interact with the HEV IRESl, further supporting its involvement in the protein synthesis process (Table 1). In addition, a literature search of IRES-binding host proteins revealed that four proteins identified in our study are known to bind other IRES. DHX9 and RPA1 bind to the IRES of HCV and FMDV; RPS3A associates with the HCV IRES, while RPL26 interacts with the p53 IRES (Table 1, highlighted in bold font) (37, 38, 39).

### Interrogation of the interaction between the HEV IRESl RNA and the host ribosomal proteins by *in silico* analyses

*In silico* molecular docking has been used in determining the complex structures of RNA-proteins interactions (40). We used HDOCK, a web-based tool which is a hybrid docking algorithm of template-based modeling for determining the interaction between the HEV IRESl RNA and four of the identified proteins (RPL5, RPL 26, RPS3A and RPS7) (41). HEV IRESl showed direct interaction with RPL5, RPL26, RPS3A and RPS7 proteins (Fig. 5A-5D). All four complexes were analyzed for identification of the hot-spot residues and was compared based on interaction mapping and binding energy analysis. All four complexes were quantified in terms of their interaction by mapping the critical polar contacts. The key nucleotides of each complex were mapped and, notably, RPS3A, RPL26 and RPL5 were found to interact with the HEV IRESl through a combination of common and unique nucleotides. The nucleotides U60, U29, G30 and U59 were common in at least two complexes. The unique nucleotides which contribute significantly in RPL5 are G50 and C44; U22, G51 and C68 in RPS3A; A38, C41 and U66 in RPL26; U9, C10, G12, G53 and G56 in RPS7 (Fig. 5E-5H). We also quantified the overall contributions by measuring the surface contacts between the HEV IRESI nucleotides and proteins (Fig. S3C). Here, the main focus was to identify the common nucleotides, which were involved in contacts with the amino acids of the different ribosomal proteins. Interestingly, we found that nucleotides A8, U9, C28, G51, G52, U59, U60, G61, U67, C68 and C69 were common in at least three proteins. Nucleotides U29, G30, G31, A38, C41, A42, G50, C55, A57, U62 and A85 were common among two proteins (Fig. S3C). Among the identified nucleotides, some are engaged with two or more amino acids such as C10, U22, U29, G30, U59 and U60, indicating their importance in mediating the interaction of the HEV IRESl RNA with the respective ribosomal protein (Fig. 5E-5H). Further investigation of the binding energy of each complex, the order of binding affinity between the four complexes are RPL5 (−391.48 kcal/mol), RPL26 (−363.43 kcal/mol), RPS3A (−341.19 kcal/mol) and RPS7 (−302.44 kcal/mol) (Fig. 5I). Physico-chemical analysis of the interacting amino acids, it was observed that most of the interacting residues were either basic (K/R) or aromatic (Y/F/H) in nature and each complex involves at least one serine residue.

**Figure 5.**
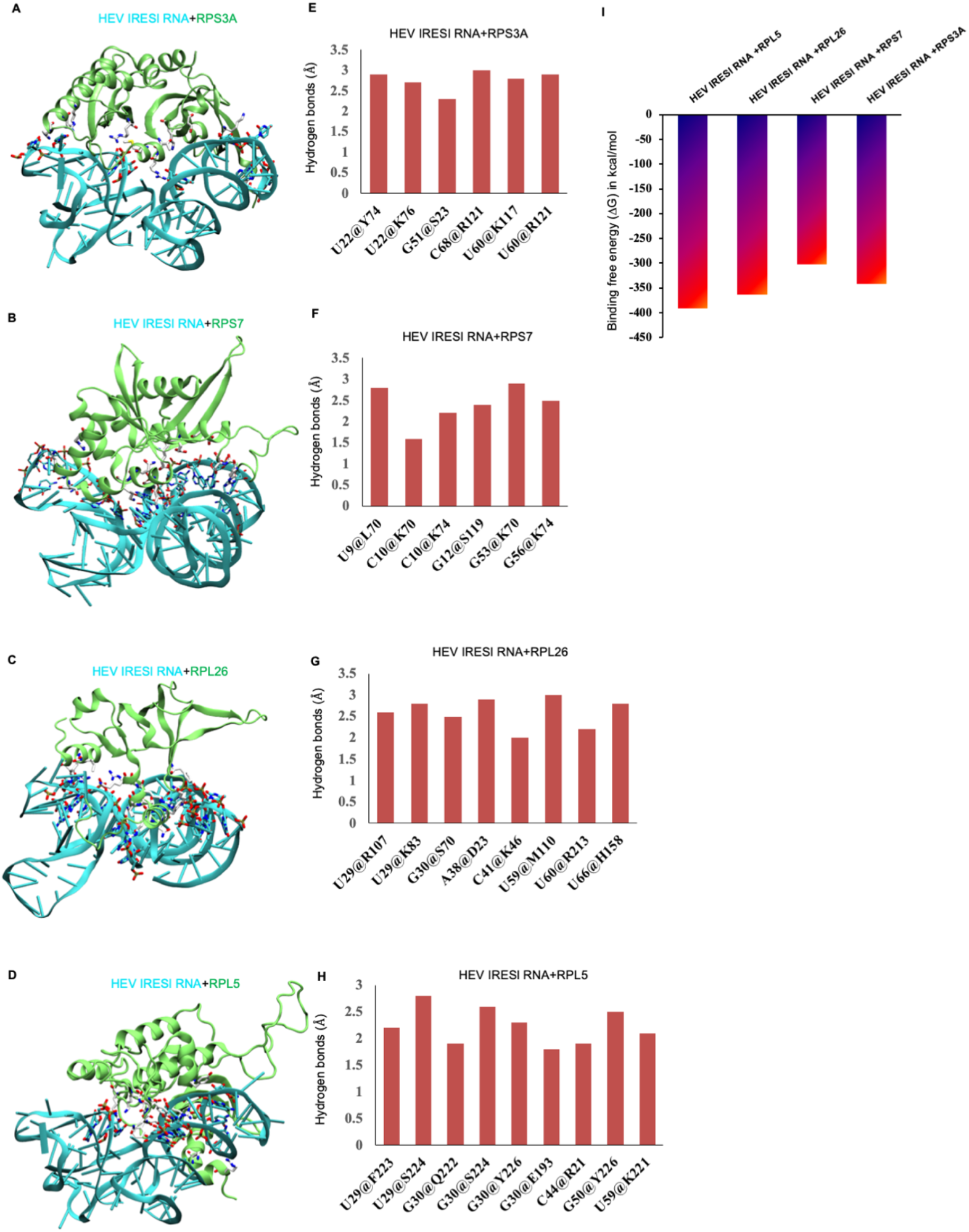
*In silico* mapping of the interaction between the HEV IRESl RNA and the ribosomal proteins. (A) The interaction between the HEV IRESI and RPS3A is shown in the cartoon and rendered in cyan and lime, respectively. The interacting nucleotides and amino acids are rendered in licorice and colored in atom-wise mode like C: cyan/white, O: red, N: blue and S: yellow. (B) The interaction between the HEV IRESI and RPS7 is shown in the cartoon and rendered in cyan and lime, respectively. The interacting nucleotides and amino acids are rendered as shown in (A). (C) The interaction between the HEV IRESI and RPL26 is shown in the cartoon and rendered in cyan and lime, respectively. The interacting nucleotides and amino acids are rendered as shown in (A). (D) The interaction between the HEV IRESI and RPL5 is shown in the cartoon and rendered in cyan and lime, respectively. The interacting nucleotides and amino acids are rendered as shown in (A). (E-H) The quantitative polar contacts between nucleotide and amino acids are measured for the indicated complexes and shown in bar-graph (Y-axis: hydrogen bond distance, X-axis interacting pair of nucleotides@aminoacids). (I) The binding free energy for the indicated complexes are plotted in the bar graph.

### RPL5 and DHX9 (RNA helicase A) proteins are essential for the function of the HEV IRESl

In order to further examine the functional significance of host interaction partners of the HEV IRESl, we used siRNA against some of the host factors to deplete them in the Huh7 cells, followed by measurement of HEV IRESl activity and g1-HEV replication. 72 hours treatment of Huh7 cells with the siRNAs against RPL5, RPL24, RPL26, RPL41, RPS3A and DHX9 depleted the corresponding proteins (Fig. 6A). Note that in the case of RPS3A siRNA, there was a significant reduction in the GAPDH protein level at 72 hours treatment period (Fig. 6A). Effect of siRNA treatment on viability of the Huh7 cells was measured at 72 hours post treatment. RPS3A siRNA treatment led to a significant loss of cell viability (Fig. 6B). Therefore, RPS3A siRNA was excluded from further studies.

**Figure 6.**
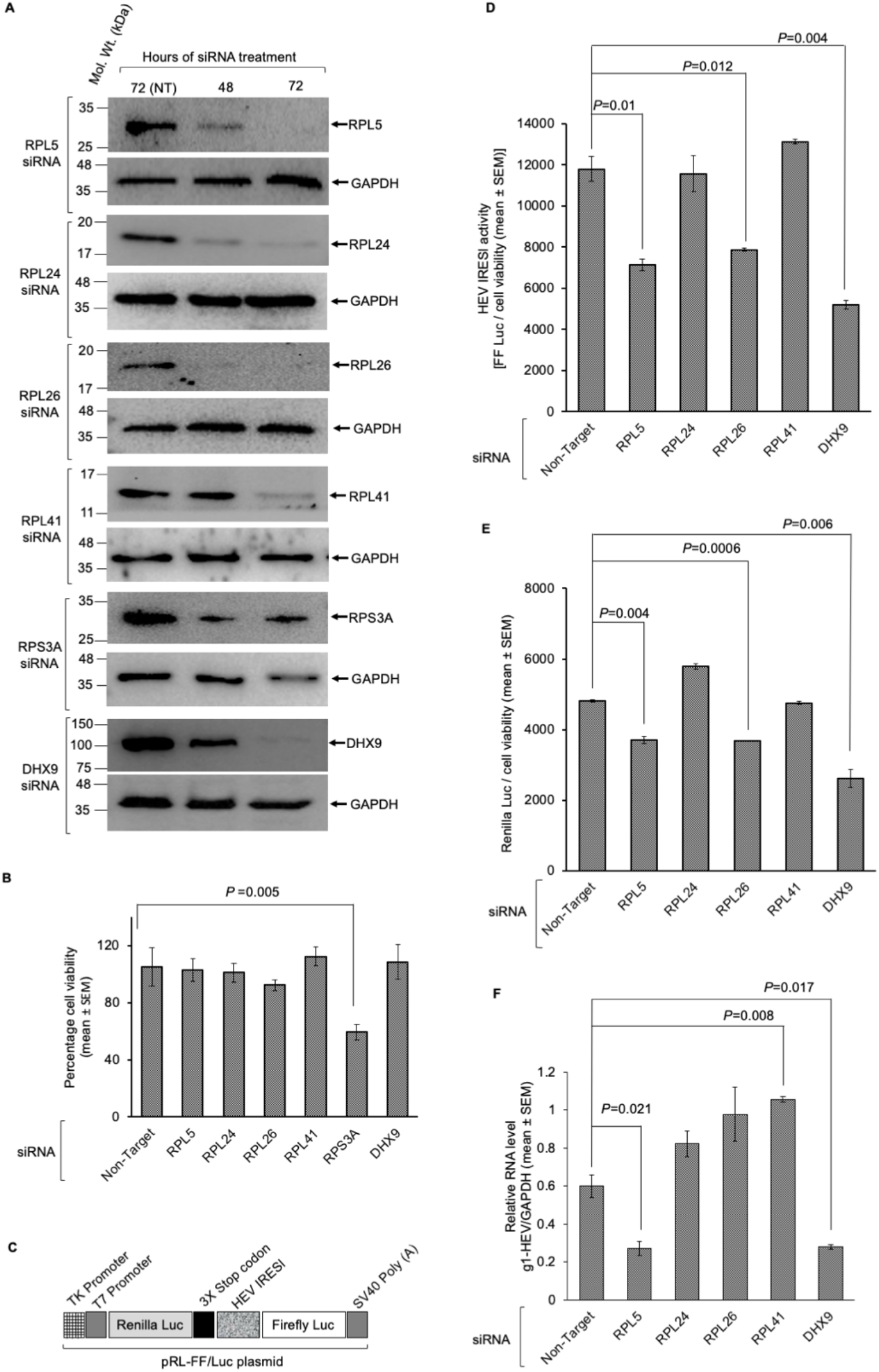
Essential roles of the ribosomal protein RPL5 and RNA helicase DHX9 in mediating the function of the HEV IRESl element. (A) Western blot analysis of the indicated proteins after 48- and 72-hours treatment with the indicated siRNAs (lanes 2 and 3). Lane 1 shows protein from cells treated for 72 hours with a non-target siRNA (NT) (B) Percent viability of Huh7 cells treated with the indicated siRNAs for 72 hours. The value for the non-target siRNA treated cells was considered to be 100% and other values were calculated relative to that. Values are the mean (± SEM) of triplicate samples (C) Schematic of the pRL-FF/Luc bicistronic reporter plasmid (D) Firefly-Luciferase activity of Huh7 cells transfected with the indicated siRNAs and the pRL-FF/Luc plasmid for 72 hours. The Firefly-Luc values were divided by that of the cell viability and represented as mean (± SEM) of triplicate samples (E) The Renilla-Luciferase activity of the Huh7 cell lysate used in (C). The Renilla-Luc values were divided by that of the cell viability and represented as mean (± SEM) of triplicate samples (F) RT q-PCR measurement of intracellular level of the g1-HEV genomic RNA in the Huh7 cells transfected with the indicated siRNAs for 72 hours. Values for the g1-HEV RNA was normalized to that of the GAPDH RNA and represented as mean (± SEM) of triplicate samples.

Next, a bicistronic reporter construct (pRL-FF/Luc dual luciferase construct) was used to simultaneously measure the HEV IRESl activity (by measuring Firefly-Luc activity) and cap-dependent translation activity (by measuring Renilla-Luc activity) in siRNA treated Huh7 cells, as described earlier (32) (Fig. 6C). Among the ribosomal proteins, siRNAs against RPL5 and RPL26 significantly reduced the activity of both Firefly-Luc (measure HEV IRESl activity) and Renilla-Luc (measure cap-dependent translation activity) (Fig. 6D, 6E). Lack of RPL24 and RPL41 did not affect the Firefly-Luc or Renilla-Luc activity (Fig. 6D, 6E). DHX9 siRNA significantly decreased the activity of both Firefly-Luc and Renilla-Luc (Fig. 6D, 6E). Next, g1-HEV expressing Huh7 cells were treated with different siRNAs, followed by measurement of the level of viral sense strand RNA. Lack of RPL5 and DHX9 proteins significantly reduced the level of g1-HEV RNA (Fig. 6F). However, lack of RPL24 and RPL26 did not affect g1-HEV RNA level and lack of RPL41 marginally enhanced the level of g1-HEV RNA (Fig. 6F).

### HEV IRESl RNA is associated with the polysomes in translationally active cells

A ribosomal fractionation assay was performed to verify the localization of the HEV IRESl element in actively translating ribosomes (Fig. 7A). A construct expressing the Firefly-Luc reporter under control of the HEV IRESl in a cap-independent manner (pSuper-HEV IRESl) was used to distinctly identify the presence of the HEV IRESl in the actively translating ribosomes. Note that HEV IRESl RNA transcription is driven by the H1 RNA polymerase III promoter in the pSuper vector. A Renilla-Luc reporter driven by cap-dependent translation was used as a positive control for the assay (pRL-TK). HEK 293T cells were cotransfected with the pSuper-HEV IRESl and the pRL-TK plasmids. 48 hours post transfection, cells were lysed and Firefly, Renilla-Luc activity was measured. As expected, significant activity of both Firefly and Renilla-Luc was observed (Fig. 7B). In order to verify if Firefly-Luc was produced through a cap-independent process, total RNA was isolated from the cotransfected cells, followed by immunoprecipitation of capped RNAs using the anti-m^7^G cap antibody, which specifically recognizes the m^7^G in the cap structure (42). Level of the Firefly or the Renilla-Luc encoding RNA was quantified by Real-Time PCR. As expected, the Renilla-Luc RNA was detected in the anti-m^7^G cap antibody immunoprecipitated sample whereas the Firefly-Luc RNA was not detectable in aliquots of the same sample (Fig. 7C). In contrast, the Firefly-Luc RNA was detected in the unbound fraction only (Fig. 7C).

**Figure 7.**
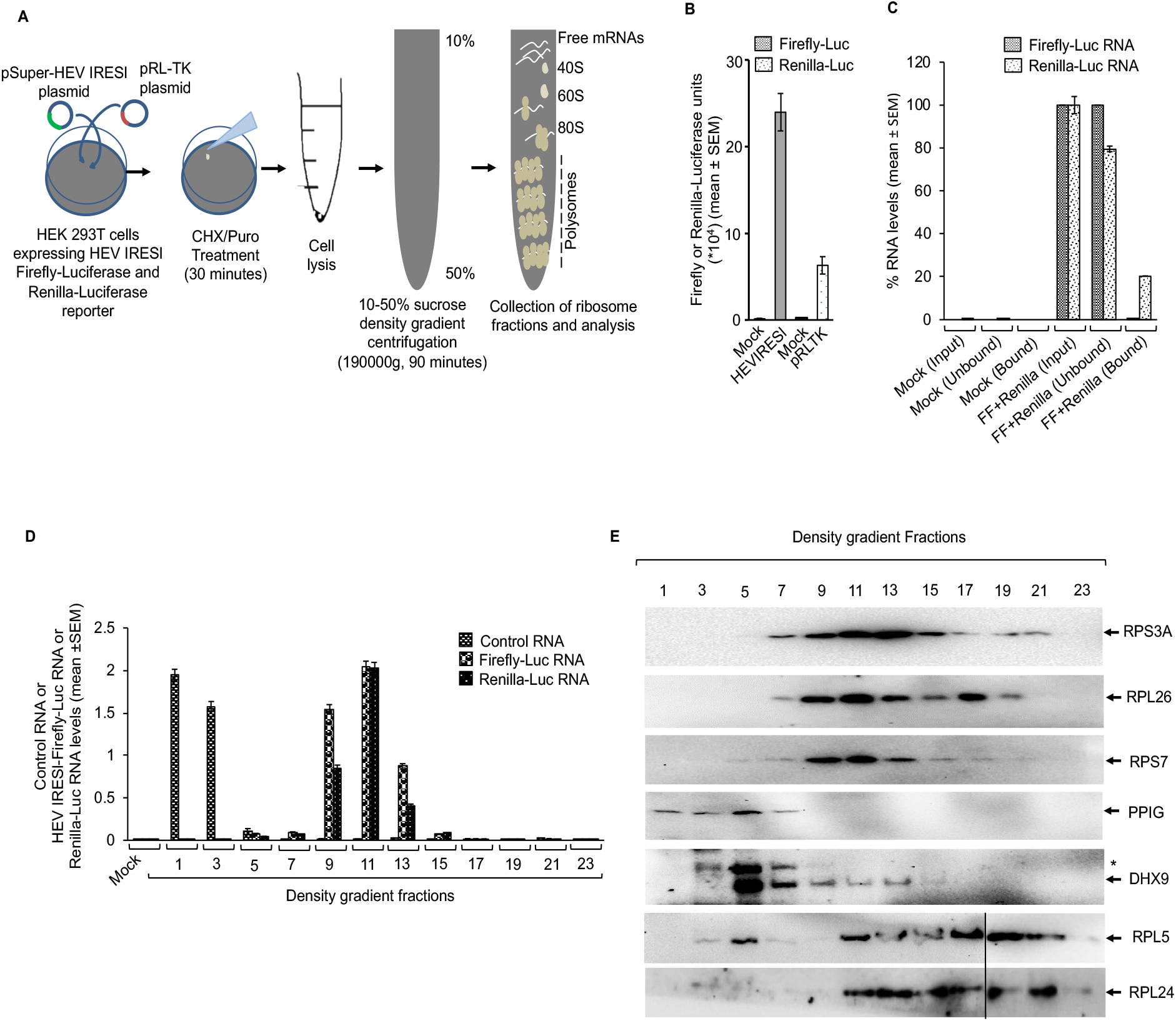
The HEV IRESl element associates with actively translating ribosomes. (A) Schematic of the ribosome fractionation assay. CHX: Cycloheximide, Puro: Puromycin (B) Measurement of the Firefly and the Renilla-Luciferase activities in the Huh7 cells, cotransfected with the pSuper-HEV IRESl and the pRL-TK plasmids. Mock shows pSuper vector transfected Huh7 cells, processed in parallel. Values are represented as mean (± SEM) of triplicate samples (C) RT-qPCR analysis of the Firefly-Luc and the Renilla-Luc RNAs obtained by the RNA-IP assay using anti-m^7^G-cap antibody. Mock: RNA from the pSuper vector transfected Huh7 cells, FF+Renilla: RNA from the Huh7 cells cotransfected with the pSuper-HEV IRESl and pRL-TK plasmids, Input: amount of RNA used for RNA-IP, Unbound: RNA isolated from supernatant of the IP samples before washing, Bound: RNA isolated from the elution fraction of the IP samples. Values are shown as mean (± SEM) of triplicate samples. (D) RT-qPCR analysis of the Control RNA, Firefly-Luc RNA and the Renilla-Luc RNA levels in aliquots of the total RNA isolated from the sucrose density gradient fractions of the Huh7 cells, cotransfected with the pSuper-control RNA plasmid, pSuper-HEV IRESl plasmid and pRL-TK plasmid and treated with cycloheximide. Values are mean (± SEM) of triplicate samples. (E) Western blot analysis of the level of indicated proteins present in the density gradient fractions, as shown in (D). “*”: non-specific band detected by the anti-DHX9 antibody.

Next, cytoplasmic fractions of HEK 293T cells cotransfected with the pSuper-HEV IRESl and the pRL-TK plasmids were subjected to linear (10-50%) sucrose density gradient centrifugation, followed by measurement of both Firefly and Renilla-Luc RNAs in different fractions. Cells were lysed after treatment with cycloheximide (CHX) or puromycin (Puro), which stalls translating ribosomes or prevents the polysome assembly, respectively (43, 44). Ribosome profiles of cycloheximide and puromycin treated cells showed an expected pattern, in agreement with the earlier studies (Fig. S4A) (44). Total RNA was isolated from aliquots of the alternate fractions from the density gradient samples and resolved by formaldehyde-agarose gel electrophoresis, followed by visualization of 28S and 18S RNA by ethidium bromide staining. As expected, 28S and 18S RNA was majorly present in the fractions corresponding to the polysomes in the cycloheximide treated samples (Fig. S4B, upper panel), which was significantly reduced in the puromycin treated samples (Fig. S4B, lower panel). RT-qPCR revealed that in cycloheximide treated cells both Firefly and Renilla-Luc RNA were predominantly detected in the fractions 9 onwards, indicating their association with the polysomes (Fig. 7D). On the other hand, control RNA was mostly present in fractions 1 and 3 (representing free RNA), indicating its inability to associate with the polysome, which further confirms the specificity of the ribosome fractionation assay (Fig. 7D). Next, aliquots of the density gradient fractions were used for western blot analysis of the different ribosomal proteins, DHX9 and PPIG. In cycloheximide treated cells, RPS3A, RPL26, RPS7, RPL5 and RPL24 were predominantly present in the polysome fractions (Fig. 7E). DHX9 was found in both 40S-80S ribosome-containing fractions and polysome fractions (Fig. 7E). PPIG was predominately found in the free mRNA or 40S-80S ribosome complex-containing fractions (Fig. 7E). PPIG was used as a control to monitor purity of the polysome fractions as it is not known to be present in the polysomes. Note that PPIG is a peptidyl-prolyl isomerase of the cyclophilin family, which catalyze the conversion between *cis* and *trans* isomers of proline (45). Collectively, these findings confirm that the HEV IRESl RNA is complexed with actively translating ribosomes.

Importance of the HEV IRESl RNA-binding translation-related proteins in driving the association of the HEV IRESl RNA with the polysomes was further assessed by a ribosome fractionation assay in Firefly and Renilla-Luc expressing, different siRNA and cycloheximide-treated HEK 293T cells. Fractions 1, 5, 9 and 13 from the density gradient centrifugation samples were analyzed to quantify the levels of Firefly and Renilla-Luc RNAs as free RNA (fraction 1), in 40S-60S ribosome complex (fraction 5) or the polysome complex (fractions 9 and 13). Treatment of cells with siRNAs against RPL5, RPL26 and DHX9 significantly reduced the HEV IRESl RNA association with the polysomes whereas siRNAs against RPL24 and RPL41 did not affect that (Fig. 8A). However, in the case of Renilla-Luc RNA, in addition to RPL5 and RPL26, treatment with RPL24 siRNA also significantly reduced the association of the Renilla-Luc RNA with the polysomes. Further, treatment with DHX9 or RPL41 siRNAs did not affect the association of the Renilla-Luc RNA with the polysomes (Fig. 8B). Next, aliquots of the above samples were analyzed for detection of the RPL5 protein (an indicator of polysome-containing fractions). As expected, RPL5 was predominately detected in the polysome-containing fractions (Fig. 8C, non-Target siRNA). Treatment with RPL5 siRNA significantly reduced its level. Treatment with RPL24 siRNA also decreased the level of polysomes whereas RPL26, RPL41 and DHX9 siRNA treatment did not affect the polysomes (Fig. 8C). Collectively, these findings confirm the association of the HEV IRESl RNA with actively translating ribosomes and suggest that RPL5 and DHX-9 help in HEV IRESl-mediated translation by holding the RNA in the polysomes whereas DHX9 is dispensable for holding the Renilla-Luc RNA (cap-dependent translation) in the polysomes.

**Figure 8.**
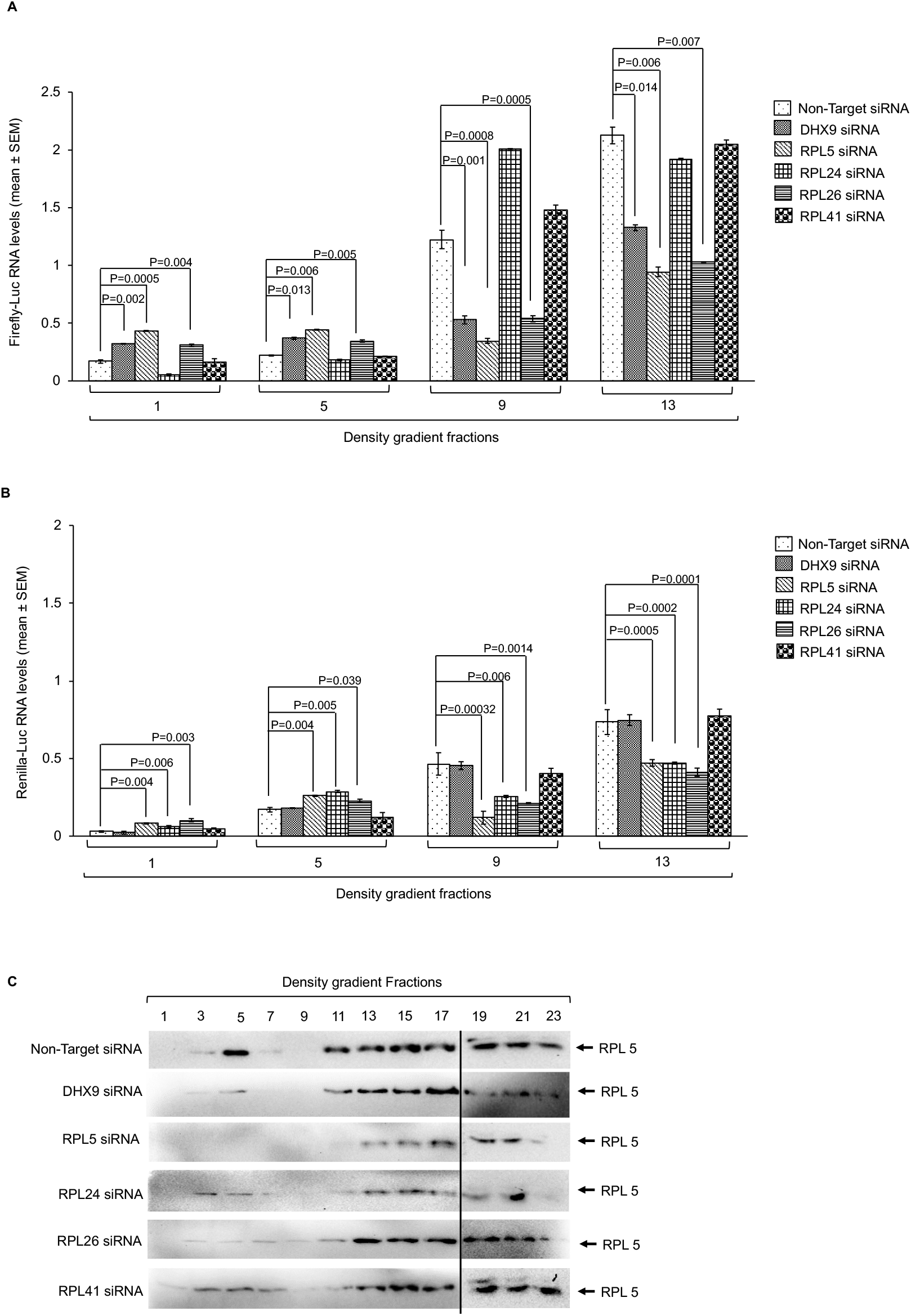
RPL5, RPL26 and RNA helicase A are important for association of the HEV IRESl element with the polysome. (A) RT-qPCR analysis of the Firefly-Luc RNA level in the density gradient fractions 1, 5, 9 and 13 of the Huh7 cells, transfected with the indicated siRNAs and the pSuper-HEV IRESl and pRL-TK plasmids for 72 hours and treated with Cycloheximide for 30 minutes. Values are mean (± SEM) of triplicate samples. (B) RT-qPCR analysis of the Renilla-Luc RNA level in the density gradient fractions 1, 5, 9 and 13 of the Huh7 cells, transfected with the indicated siRNAs and the pSuper-HEV IRESl and the pRL-TK plasmids for 72 hours and treated with Cycloheximide for 30 minutes. Values are mean (± SEM) of triplicate samples. (C) Western blot analysis of the RPL5 protein level in the indicated density gradient fractions of the Huh7 cells, transfected with the indicated siRNAs and the pSuper-HEV IRESl and the pRL-TK plasmids for 72 hours and treated with Cycloheximide for 30 minutes.

## Discussion

While investigating the cause of poor replication of the g1-HEV in the Huh7 cells, we observed better viral replication upon treatment of cells with ER stress-inducing compounds such as tunicamycin and thapsigargin. Subsequently, we identified a fourth ORF in the viral genome, which produced the ORF4 protein that played a key role in the assembly of the viral replication complex (32). We also showed that ORF4 was synthesized by an IRES-like (HEV IRESl) element. Sequence analysis of the HEV IRESl indicated its weak homology with the Equine Rhinitis A virus (ERAV) IRES (a type II IRES) (32, 46). The HEV IRESl element is remarkably smaller in size than other canonical IRES elements and it requires only 87 nucleotides to drive the cap-independent translation of the ORF4 protein in its native context as well as to drive the translation of a heterologous protein such as the Firefly-Luciferase, when inserted into a bicistronic reporter construct. In contrast, function of known IRES elements depend on double the number of nucleotides or more. Smaller size and relatively simple secondary structure of the HEV IRESl element offers the advantage of dissecting out the molecular mechanisms that are crucial for initiation of cap-independent internal translation. Therefore, the current study was undertaken to further substantiate the claim that HEV IRESl element is a bonafide internal ribosome entry site.

Two independent approaches were followed to identify the host interaction partners of the HEV IRESl RNA: RaPID assay and Y3H assay. The RaPID assay identifies stable and transient interactions of all proteins associated with the RNA of interest that includes both direct and indirect interaction partners of the RNA. Several controls were used to avoid/subtract non-specific interaction partners, thus ensuring generation of high confidence mass spectrometry data for downstream studies. For example, good quality of our HEV IRESl RaPID assay-LC-MS-MS data is evident from the fact that comparison of the HEV IRESl-interacting proteins with the SARS-CoV2 5’-and 3’-UTR RNA-interacting proteins (reported by our laboratory using the same technique) identified only RPL7a as the common protein (34). RPL7a is an essential component of the large subunit of ribosome and known to directly bind with RNA through its two distinct RNA binding domains (47). Presence of RPL7a in both the datasets seems to be in agreement with its required function rather than being an experimental artefact.

A limitation of the RaPID assay is attributed to its dependence on biotinylation of lysines exposed on the surface of RNA-binding proteins present within 10 nm distance from the BirA ligase. Thus, it is possible to lose some of the RNA binding proteins if they are buried deep inside the complex and inaccessible to BirA ligase for biotinylation. Therefore, a conventional Y3H assay-based human liver cDNA library screening was conducted in parallel to identify the interaction partners of the HEV IRESl RNA. Y3H assay offers the advantage of identifying the direct interaction partners of a bait RNA. Y3H assay identified ribosomal proteins (RPL5, RPL26, RPL41, RPS7, RPS15A and RPS3A), a known DNA binding protein (GREBP) and a cyclophilin family protein (PPIG) as interaction partners of the HEV IRESl (48, 45). Specificity of the Y3H data was further ensured by assessing the interaction of the eight HEV IRESl RNA-interacting proteins with two well-known IRES sequences such as the FMDV IRES (Type II IRES) and the HCV IRES (Type IV IRES). Distinct patterns of interaction were observed for these RNA sequences, supporting the specificity of the interactions detected in Y3H assay. Although both RaPID and Y3H assays identified specific interaction partners of HEV IRESl, no common proteins were detected in the two assays. A search of the initial RaPID data of HEV IRESl sample identified RPL26 as a common protein present in both RaPID and Y3H data sets. However, RPL26 was not present in the final list of HEV IRESl interaction partners as it had a PEP score and Prot score of 7 and 29, respectively, whereas the cutoff threshold was set to 17 and 30, respectively. Note that RPL26 was not present in the negative control sample RaPID data.

In order to further confirm the interaction between the HEV IRESl RNA and the host proteins, *in vitro* biotinylated-RNA pull down assay and molecular docking analysis was performed for some of the HEV IRESl interacting proteins, which supported their interaction. Docking analysis also demonstrated that the complex between the HEV IRESl and RPL5 was the most stable with respect to the others. Moreover, comparative interaction mapping showed that the interaction of the HEV IRESl element with the ribosomal proteins majorly involved the SL A (nucleotides U9, C10, G12, U22), SL B (nucleotides U29, G30, A38, C41, C44, G50) and the unpaired region between the SL B and SL C (G51, G53, G56, U59, U60, U66, C68) of the former (refer to the HEV IRESl secondary structure in Fig. 1A). In summary, multiple validation steps ensured that the HEV IRESl RNA interaction partners identified in our experiments are of high confidence.

Bioinformatics analysis of the 51 HEV IRESl RNA-binding proteins revealed enrichment of proteins involved in the translation process. Eight ribosomal proteins and six translation regulatory factors interact with the HEV IRESl RNA, in support of its function as a translation initiation site. Four proteins identified in our study are known to interact with the IRES of other viruses.

Silencing of RPL5 significantly inhibited the HEV IRESl activity as well as the g1-HEV replication. RPL5 strongly interacts with the HEV IRESl, weakly interacts with HCV IRES and does not interact with FMDV IRES in Y3H assay. RPL5 showed lowest binding free energy (ΔG) for interaction with the HEV IRESl RNA, *in silico*. RPL5 is also a component of the polysome and lack of RPL5 significantly affected the polysome stability and dissociated both the HEV IRESl and the Renilla-Luc RNAs from the polysome. Hence, it is clear that RPL5 is an essential factor for both cap-dependent and cap-independent translation process. RPL26 showed a pattern similar to RPL5 by inhibiting the HEV IRESl-Luc and Renilla-Luc activities. Moreover, the HEV IRESl-Luc and the Renilla-Luc RNA association with the polysome was significantly reduced upon treatment of cells with the RPL26 siRNA. However, RPL26 differed from RPL5 in the following parameters: it bound moderately with the HEV IRESl RNA both in Y3H assay and *in silico* analysis, lack of RPL26 did not affect g1-HEV replication and lack of RPL 26 did not affect the polysome stability. Earlier studies have shown that lack of RPL26 does not affect cap-dependent translation in *Saccharomyces cerevisiae* and RPL26 is a known ITAF, which has been shown to control DNA damage induced cap-independent translation of the p53 mRNA (49, 50, 51). Based on our data and available literature, we propose that RPL26 may be involved in mediating efficient function of the HEV IRESl though it is not an essential factor. Ablation of both RPL5 and RPL26 may show a more profound effect on the HEV IRESl activity. Further studies are required to clarify the role of RPL26 in mediating the function of HEV IRESl. Silencing of RPL24 and RPL41 did not inhibit either the HEV IRESl activity or the Renilla-Luc activity. Lack of RPL24 reduced the stability of the polysomes, which is in agreement with earlier reports (52). However, lack of RPL24 did not affect the HEV IRESl RNA association with the polysomes although it reduced the association of the Renilla-Luc RNA with the polysomes, indicating its importance in cap-dependent translation. Lack of RPL41 did not affect the polysome stability or HEV IRESl and Renilla RNA association with the polysomes. Interestingly, lack of RPL41 significantly increased the g1-HEV replication, possibly due to extra-translational function of RPL41. Note that RPL41 knockdown has been shown to arrest the cell-cycle in the G2/M phase and it acts by stabilizing the microtubule (53). Our earlier study has shown that HEV replication is significantly high in the G2/M phase of the cell-cycle (54).

Apart from the ribosomal proteins, DHX9/RNA helicase A (RHA), HNRNPA2/B1, YTHDF2, EEF2 and LARP4 were also identified as interaction partners of the HEV IRESl RNA. RHA is known to associate with the IRES of FMDV and HCV and acts as an ITAF. It also promotes translation from the IRES of retroviral RNA and jun D mRNA and regulates the replication of several RNA viruses, including the porcine reproductive and respiratory syndrome virus, bovine viral diarrhea virus, classical swine fever virus and dengue virus (55–58). It binds to the 3’-UTR of HEV; however, significance of the interaction is not known (59). RHA is a member of the DEx H-box family of superfamily II (SF2) helicases, which unwind duplex RNA possessing a 3’-single stranded RNA tail by hydrolyzing the nucleoside triphosphates (60, 61). In our experiments, lack of RHA significantly reduced both the HEV IRESl and the Renilla-Luc activities as well as reduced the g1-HEV replication. It was found to be associated with the 40S-80S ribosomes and polysomes, however, its absence did not affect the polysome stability. Lack of RHA resulted in reduced recruitment of the HEV IRESl RNA to the polysomes, however, Renilla-Luc RNA association with the polysomes was not affected. Thus, it is clear that RHA is involved in controlling the activity of the HEV IRESl. HNRNPA2B1 is reported to be involved in mRNA processing. It has been shown to bind to the 3’-UTR of CDK6 and recruit RHA, leading to miRNA mediated silencing of CDK6 (62). An earlier study has shown that HNRNPA2B1 binds to the genomic and sub genomic promoters of HEV and plays an essential role in the g1-HEV replication. Further, intracellular localization of HNRNPA2B1 is altered in the HEV infected cells (63). YTHDF2 is known to accelerate mRNA decay by targeting the mRNAs to cytoplasmic processing bodies (P-bodies) and by recruiting the CCR4-NOT deadenylase complex (64, 65). EEF2 (eukaryotic elongation factor 2) is a GTP binding translation elongation factor, which plays an essential role in translation by promoting GTP dependent translocation of the ribosome (66). LARP4 binds to the poly-A tract of mRNA, associates with the 40S ribosomal subunit as well as polysomes and plays a role in translation regulation by preventing deadenylation at poly-A tail (67–69). Presence of these factors in the HEV IRESl RNA-associated protein complex suggests their potential role(s) in regulating the HEV IRESl-mediated translation. Further studies are warranted to clarify their mode of action.

An important aspect of the HEV IRESl function pertains to upregulation of its activity by the ER stress inducing agents in the g1-HEV infected cells (32). The RaPID assay identified four proteins involved in the ER stress response pathway such as the heat shock protein 105 kDa (HSP 105), the luminal ER protein of 29 kDa (ERp29), ψ-taxilin and Pumilio homolog 1 (PUM1) as interaction partners of the HEV IRESl. HSP 105 is known to regulate the ER stress induced caspase-3 activation (70). ERP29 is induced by the ER stress. It is ubiquitously expressed and implicated in the biosynthesis and trafficking of many proteins (71). γ-taxilin belongs to the syntaxin-binding protein family, it interacts with ATF4 and regulates the ER stress response (72). PUM1 is known to be phosphorylated by the IRE1 kinase under conditions of ER stress, it binds spliced XBP1 RNA and prevents its degradation (73). Upon ER stress, interaction between PUM1 and HEV IRESl might be protecting the later from degradation by the regulated IRE1-dependent decay (RIDD) pathway.

In conclusion, we have identified 51 host proteins, which associate with the HEV IRESl RNA in mammalian cells. Four of the identified proteins have been reported to associate with known IRES elements of viral and cellular origin and two of the identified proteins are known ITAFs. Several ribosomal proteins, translation regulatory factors, components of the tRNA synthetase complex were found to be associated with the HEV IRESl RNA. Taken together, above data unequivocally demonstrate that the HEV IRESl is a bonafide internal ribosome entry site capable of driving cap-independent translation. These findings significantly advance our understanding of the molecular mechanisms that control the synthesis of the ORF4 protein in the g1-HEV infected cells. Moreover, considering the small size and simplicity of the secondary structure of the HEV IRESl element compared to other well-known IRES elements, it is a useful tool for recombinant protein expression as well as for investigating the molecular details of the cap-independent IRES-mediated translation process.

## Materials and Methods

### Plasmids and reagents

The RNA motif plasmid cloning backbone [(pRMB) (Addgene plasmid # 107253; http://n2t.net/addgene:107253; RRID: Addgene_107253)] and the BASU RaPID plasmid (Addgene plasmid # 107250; http://n2t.net/addgene:107250; RRID:Addgene_107250) were gifted by Prof. Paul A. Khavari (School of Medicine, Stanford University, Clifornia, USA) and reported earlier (29). To clone the HEV IRESl in the pRMB vector, 2701 to 2787 nucleotides of the g1-HEV genome (Genbank ID: AF444002.1) was PCR amplified from the pSK HEV2 plasmid (32) with the following primers: IRESl87 BASU CBB FP (5’ GCTTGGTGCCGATCGGTCCC 3’) and IRESl87 BASU CBB RP (5’AGCTCATCTGGCAGCAAGCTCAG 3’) and ligated with the pRMB vector backbone generated by BSMB1 restriction enzyme digestion, followed by blunting. For hybrid RNA generation for Y3H assay, HEV IRESl, FMDV IRES and HCV IRES sequences were PCR amplified from the pSKHEV2 plasmid, pVitro2-neo-mcs plasmid (InvivoGen, San Diego, USA) and S52/SG-Feo(AI) replicon [gifted by Prof. Charles M. Rice, (74)], respectively, using the following primers: HEV IRESl FP: (5’ ACTATCTCGAGGGTGCCGATCGGTCCC 3’), HEV IRESl RP: (5’ ACTATCCATGGCATCTGGCAGCAAGCTCAG 3’); FMDV IRES FP: (5’ ACTATCTCGAGAGCAGGTTTCCCCAATGACACA 3’), FMDV IRES RP: (5’ ACTATCCATGGAAAGGAAAGGTGCCGACCTCCG 3’); HCV IRES FP: (5’ ACTATCTCGAGACCTGCCTCTTACGAGGC 3’) and HCV IRES RP: (5’ ACTATCCATGGTTCTTTGAGGTTTAGGAAGTGTGCT 3’). PCR products were digested with the Xho1 restriction enzyme and ligated with the XhoI and SmaI digested p3HR2 vector backbone. To clone the negative sense HEV IRESl (HEVIRESl^-^) fragment in the p3HR2 vector, 87 nucleotides of IRESl sequence was PCR amplified from the pSK HEV plasmid using HEV IRESl FP and HEV IRESl RP primers and ligated with the SmaI digested p3HR2 vector. To clone the HEV IRESl sequence in the pSuper vector, pRLFF/Luc87 and pSuper plasmids were restriction digested with the BamHI and KpnI enzymes, respectively, followed by treatment with Klenow (end-filled by Klenow) and digestion of both plasmids with XhoI and ligation. To clone the control RNA (618-738 nt in g1-HEV genome) into pGL3 vector (for use in ribosome fractionation assay), pGL3 vector was digested with HindIII and NcoI followed by blunting to generate the vector backbone. Next, control RNA sequence was amplified by PCR from pSKHEV2 (Genbank ID: AF444002.1) using Control RNA FP: 5’ GCCCCCTGGCACATA 3’ and Control RNA RP: 5’ GAGCGCAGGTTGGAAA 3’. Linearized pGL3 vector backbone and control RNA sequence insert was ligated and positive clones were selected by restriction digestion. For biotinylated RNA synthesis, HEV IRESl and control RNA sequences were PCR amplified using primers as described above and cloned into pJET1.2 vector, following manufacturer’s instruction (Thermo Scientific, Massachusetts, USA). All clones were confirmed by DNA sequencing. All plasmids are available upon request.

Human liver cDNA library was obtained from Clontech (California, USA). Dual Luciferase reporter assay kit and CellTiter96 Aqueous one solution cell proliferation assay kits were from Promega (Madison, USA). Non-targeting siRNA (Catalog no. D-001810-10-20), Human RPL5 (6125) siRNA (Catalog no. L-013611-00-0005), Human RPL26 (6154) siRNA (Catalog no. L-011132-01-0005), Human RPL24 (6152) siRNA (Catalog no. L-011144-02-0005), Human RPL41 (6171) siRNA (Catalog no. L-011160-01-0005), Human RPS3A (6188) siRNA (Catalog no. L-013607-00-0005) Human DHX9 (1660) siRNA (Catalog no. L-009950-00-0005) were from Ge Healthcare Dharmacon (Colorado, USA). Antibodies against RPL5 (Catalog no. A303-933A) and RPL26 (Catalog no. A300-686A) were from Bethyl laboratories (Texas, USA). Anti-RPL24 (Catalog No. ITT09128), anti-RPL41 (Catalog No. ITT09136) and anti-DHX9 (Catalog No. ITT12266) antibodies were from Geno Technology Inc. (St Louis, USA); anti-RPS3A (catalog No. STJ28158), anti-RPS7 (catalog No. STJ28814) and anti-PPIG (catalog No. STJ191133) antibodies were from St. John’s laboratory (London, UK). Anti-GAPDH (catalog no. sc-25778) antibody was from Santa Cruz Biotechnology (Texas, USA). Goat anti-rabbit IgG-HRP (Cat No. 4030-05) was from Southern Biotech (Alabama, USA). 3-Amino 1, 2, 4 triazole (3-AT), Ortho-Nitrophenyl-β-galactoside (ONPG), Cycloheximide and Puromycin was from Sigma (Missouri, USA). Cycloheximide and puromycin were used at a working concentration of 100μg/ml and 0.5μg/ml, respectively.

### Mammalian cell-culture, transfection, *in vitro* transcription and cell viability assay

HEK 293T cells were obtained from ATCC (Virginia, USA). Huh7 cells were as described previously (32). Cells were maintained in Dulbecco’s modified Eagle medium (DMEM) containing 10% Fetal Bovine Serum (FBS), 50 I.U./mL Penicillin and Streptomycin at 37^0^C in a 5% CO_2_ incubator. Cells were maintained in antibiotic free medium before starting the experiments. For plasmid transfection, cells were seeded at 70-80% confluency in DMEM + 10% FBS and incubated overnight at 37^0^C in a 5% CO_2_ incubator. Next day, cells were transfected with the desired plasmids using Lipofectamine 2000 transfection reagent (Thermo Scientific, Massachusetts, USA), at 1:1 ratio, following manufacturer’s instruction. 6-8 hours post-transfection, culture medium was replaced with fresh DMEM+10% FBS.

For experiments involving siRNA mediated gene silencing, cells were seeded at 70-80% confluency on 12 well TC dishes and incubated overnight at 37^0^C with 5% CO2. Next day, 25nmol siRNA was transfected into each well using 0.35μl Dharmafect transfection reagent, following manufacturer’s instruction (GE Healthcare Dharmacon Inc., Colorado, USA). 12 hours post-transfection, culture medium was replaced with fresh medium (DMEM + 10% FBS) and cells were maintained at 37^0^C in a 5% CO_2_ incubator.

The HEV genomic RNA was *in vitro* synthesized as capped RNA, using mMessage mMachine kit (Thermo Scientific, Massachusetts, USA), as described (32). Size and integrity of the *in vitro* synthesized RNA was monitored by formaldehyde agarose gel electrophoresis. *In vitro* synthesized RNA was transfected using Lipofectamine 2000 transfection reagent, at 1:1 ratio, following manufacturer’s instruction (Thermo Scientific, Massachusetts, USA). 6-8 hours post-transfection, the culture medium was replaced with fresh medium (DMEM+10% FBS) and maintained until further manipulation.

For biotinylated-RNA pull down assay, pJET1.2 HEV-IRESl and pJET1.2 Control RNA plasmids were linearized by restriction digestion with NcoI enzyme and column purified. One microgram of purified linearized DNA was used for *in vitro* transcription reaction using Maxi-script T7 transcription kit, following manufacturer’s instruction (Thermo Scientific, Massachusetts, USA). 0.2mM Biotin-16-UTP (Sigma-Aldrich, REF:11388908910) and 0.3mM unlabeled-UTP were added to the reaction mixture to produce biotinylated RNA. Non-biotinylated RNAs were transcribed in parallel in the presence of 0.5mM unlabeled UTP. *In vitro* transcribed RNA was purified as per the instruction of the manufacturer of Maxi-script kit.

Cell viability was measured using a commercially available kit [CellTiter 96 Aqueous One Solution Cell Proliferation Assay (Promega, Madison, USA)], which utilizes Tetrazolium salt based colorimetric assay. Details are as described (32).

### Yeast three Hybrid (Y3H) Assay and screening of the human liver cDNA library

A GAL4 based Y3H assay system, gifted by Prof. Marvin Wickens (University of Wisconsin, Madison, USA) was used to determine the RNA-protein interactions and identification of the HEV IRESl RNA-binding proteins by screening of the human liver cDNA library using the YBZ1 Yeast strain (35).

The YBZ1 competent cells were prepared following Lithium Acetate method, as described (75). To examine self-activation by the p3HR2-HEV IRESl, 2µg each of the p3HR2-HEV IRESl and pACT2 plasmids or p3HR2 and pACT2 plasmids were cotransformed into the YBZ1 competent cells and plated on LU^-^ media, as described (75). After 3 days of incubation in a humidified incubator at 30^0^C, 8 random colonies were replica plated onto LUH^-^ as well as 3-AT (0.1-1 mM) supplemented LUH^-^ plates, followed by 4 days incubation at 30^0^C in a humidified incubator. Colonies were monitored on the 4^th^ day. The p3HR2-HEV IRESl-containing colonies showed very little/no growth on the LUH^-^ and LUH^-^ +3-AT media (Figure S2).

Next, 2µg of the p3HR2-HEV IRESl plasmid DNA was transformed into the YBZ1 competent cells and plated on the U^-^ media. After 3 days of incubation at 30^0^C, a single colony was inoculated in the U^-^ media and grown at 30^0^C to prepare the primary culture (YBZ1-p3HR2-HEV IRESl) and subsequently, the secondary culture. Aliquots of the culture were used for RNA isolation and verification of the HEV-IRESl RNA expression by RT-PCR. Next, the secondary culture grown from the primary culture was used for preparation of competent cells of the YBZ1-p3HR2-HEV IRESl.

The human liver Matchmaker cDNA library was procured in the *E.coli BNN132* strain with a titer of 210^8^ cfu/ml. The cDNA library contained 3.5*10^6^ independent clones, with ∼93% colonies containing the cDNA insert. The cDNA library was amplified and a stock titer of 210^7^ cfu/ml was prepared and stored at −80^0^C until further use, following the manufacturer’s instruction. Plasmid DNA was isolated from the retitered stock and used for transformation into the YBZ1-p3HR2-HEV IRESl competent cells (YBZ1 strain containing the p3HR2-HEV IRESl plasmid). A pilot transformation was performed to estimate the efficiency of the cDNA library and optimize the ideal quantity of DNA required for screening, following the instructions of the manufacturer. Approximately, 4*10^6^ independent library clones were screened on the LUH^-^ + 0.5mM 3-AT media-containing plates (0.5mM 3-AT was added to enhance the stringency of the screening), leading to growth of 395 colonies. Subsequent replica plating of those 395 colonies on LUH^-^ plates containing an increasing concentration of 3-AT (2mM, 5mM, 10mM and 20mM) showed that all colonies could grow on 10mM 3-AT or more. These colonies were inoculated in 5 ml LU^-^ media and incubated overnight at 200 RPM, 30^0^C, in an incubator-shaker. Next day, cells were harvested by centrifugation at 7000×g, 4^0^C, 10 minutes. Plasmid DNA was isolated using the Qiaprep miniprep kit (Qiagen, Hilden, Germany). Briefly, cell pellet was resuspended in the buffer P1 (supplied in the kit), followed by 6 cycles of freeze-thaw in liquid nitrogen and ambient temperature, respectively. Next, the buffer P2 was added, samples were mixed thoroughly and incubated for 5 minutes at ambient temp. Next, buffer N3 was added, samples were mixed thoroughly and centrifuged at 18000×g, 4^0^C, 10 minutes. Supernatant was loaded onto the kit-supplied columns and plasmid DNA was eluted after a series of washing, as per the instruction of the manufacturer. Plasmid DNA isolated from the YBZ1 transformants were transformed into the *E.coli Top_10_* strain and colonies were grown on the LB agar media containing ampicillin (100 µg/ml). Presence of the pACT2 plasmid with a cDNA insert sequence in the *E.coli* transformants was verified by a colony PCR using the following primers: pACT2 AD-LT seq FP: (5’ ATT CGATGATGAAGATACCCCA 3’) and pACT2 AD-LT seq RP: (5’ GTGAACTTGCGGGGTTTTTCAGTATCTACGA 3’). 285 colonies were positive in the PCR, indicating that they contained a library clone with a cDNA insert. Plasmid DNA was isolated from these colonies and digested with HindIII, XhoI and EcoRI restriction enzymes to sort them into different categories based on their restriction patterns. 75 clones were found to contain unique cDNA insert. The insert and its flanking region was sequenced using the following primer: pACT2 AD LT SEQ Primer: TGGTGGGGTATCTTCATCATCGAATAG. Analysis of the sequencing data identified 8 unique protein coding gene sequences in frame with the GAL4 AD. These clones were cotransformed along with the p3HR2-HEV IRESl plasmid into the YBZ1 competent cells with appropriate controls to ensure that the interaction is reproducible and there is no self-activation by the prey protein. Liquid β-galactosidase assay of the colonies were done as described earlier (75).

### RaPID assay

The RaPID assay was done as described, following similar conditions (34). To identify the HEV IRESl-interacting host proteins, HEK 293T cells were co-transfected with the pRMB HEV IRESl and BASU plasmids (in 6:1 ratio) using lipofectamine 2000 (1:1 ratio). 42 hours post-transfection, 200µM biotin supplemented media (DMEM+10%FBS) was added to the cells and maintained for 18 hours.

#### Cell lysate preparation

Cells were washed thrice with ice-cold PBS and lysed in prechilled radio-immunoprecipitation (RIPA) buffer (150 mM NaCl, 1% NP-40, 0.5% sodium deoxycholate, 0.1% SDS, 50mM Tris, pH 8.0), supplemented with protease and phosphatase inhibitor cocktail. The lysate was centrifuged at 14000×g for 40 min and a Macrosep advanced filter (3000 Da MW cutoff, 20 ml; catalogue no. 89131-974; VWR, USA) was used to remove the free biotin. Clarified supernatant was precipitated in acetone, first at −20°C for 10 minutes and then at −80°C for 20 minutes. Protein precipitate was solubilized in 8M urea and Bicinchoninic (BCA) protein assay kit (Thermo Fisher Scientific, MA, USA) was used to determine the protein concentrations of each sample.

#### Digestion and peptide preparation

10mg protein for each sample was treated with 10mM DTT (Dithiothreitol), for 30 min at 56^0^C and alkylated with 20mM iodoacetamide (IAA) at room temperature for 1 hour in dark. Trypsin (catalog no. T1426; Thermo Fisher Scientific, MA, USA) was added to the samples at a 1: 20 (wt/wt) ratio and incubated at 37^0^C for 24 hours. Next, 1% formic acid was added to the samples and desalting of the peptides was done using a Sep-Pak C18 cartridge (catalog no. WAR020515; Waters, MA, USA) followed by lyophilization in a SpeedVac. Lyophilized peptides were solubilized in 1ml PBS and incubated with 150µl pre-washed streptavidin agarose beads (catalog no. 20361; Thermo Fisher Scientific, MA, USA) for 2 hours at ambient temperature. The beads were washed in PBS, followed by washing in wash buffer (5% acetonitrile in PBS) and finally with ultrapure water. Excess liquid was completely removed from beads, and biotinylated peptides were eluted by adding 0.3 ml of a solution containing 0.1% formic acid and 80% acetonitrile in water by boiling at 95°C for 5 min. A total of 10 elutions were collected and dried together in a Speed Vac. Enriched peptides were desalted with C18 tips (Thermo Fisher Scientific, MA, USA), and reconstituted with solvent A (2%, v/v, acetonitrile, 0.1% v/v, formic acid in water) for LC-MS/MS analysis.

#### LC-MS/MS acquisition

A Sciex 5600^+^ triple time of flight (TOF) mass spectrometer was used for LC-MS/MS analysis. It was coupled with a chromXP reversed-phase 3-µm C_18_-CL trap column (350 μm by 0.5 mm, 120 Å; Eksigent; AB Sciex, MA, USA) and a nanoViper C_18_ separation column (75 μm by 250 mm, 3 μm, 100 Å; Acclaim PepMap; Thermo Fisher Scientific, MA, USA) in an Eksigent nanoLC (ultra 2D plus) system. A binary mobile solvent system was applied, with the following composition: solvent C, 2% (v/v) acetonitrile, 0.1% (v/v) formic acid in water; solvent B, 98% (v/v) acetonitrile, 0.1% (v/v) formic acid. To separate the peptides, a 60 minutes gradient was run with a total run time of 90 minutes at a flow rate of 200 nl/minute and MS data was acquired in information dependent-acquisition (IDA) with high sensitivity mode. Each cycle was run for 2.3 seconds with 250- and 100-ms acquisition times for MS1 (*m/z* 350 to 1,250 Da) and MS/MS (100 to 1,800 *m/z*) scans, respectively. Every experimental condition was run in quadruplets.

#### Protein identification and quantification

All raw files (.wiff) were searched in protein Pilot software (version 4.5; Sciex) with Mascot algorithm for protein identification and semi quantitation against Swiss-Prot_57.15 database (20,266 sequences after application of *Homo sapiens* taxonomy filter). Following parameters were applied to search biotinylated peptides: (a) Trypsin as a proteolytic enzyme with maximum of two missed cleavages allowed; (b) Allowed tolerance limit for peptide mass error was 20ppm;(c) Mass error tolerance for fragments was taken up to 0.20 Da; (d) Carbamido-methylation of cysteine (+ 57.02146 Da), oxidation of methionine (+15.99491 Da), deamination of NQ (+ 0.98416), and biotinylation of lysine (+ 226.07759 Da) were taken as variable modifications. Pearson correlation plot of peptide intensity was used to monitor the quality of data between each run and replicates.

#### Data analysis

A protein was considered to be identified if it corresponds to one or more biotinylated peptide, with PEP (posterior error probability) score ≥ the median value in the Gaussian smoothing curve of each sample. Note that PEP score refers to the probability that the observed peptide spectrum matches are incorrect. A web-based tool, Bioinformatics and Evolutionary Genomics (http://bioinformatics.psb.ugent.be/ webtools/Venn/) was used to identify host proteins that are specific to HEV IRESl RNA and formation of Venn diagram. Data set with a minimum of 1 unique peptide and a “prot score” of 30 or more was considered in the final list of host proteins for further studies (Table 1). “Prot Score” refers to the overall protein score generated by Mascot, considering all the observed mass spectrums, which match amino acid sequences of a certain protein.

#### Data availability

The mass spectrometry proteomics data have been deposited to the ProteomeXchange Consortium via the PRIDE partner repository with the dataset identifier PXD031009.

### Bioinformatics analysis

RNA secondary structures were analyzed using the Mfold program (http://www.unafold.org/mfold/applications/rna-folding-form.php), based on a minimum free energy calculation at 25^0^C (76). The host-virus RPPI data set was visualized using Cytoscape (version 3.1.0). Network Analyzer plug-in in Cytoscape was used to compute the topological parameters and centrality measures of the network. Gene ontology (GO) and Reactome pathway analysis were performed using the Gene Set Enrichment (https://www.gseamsigdb.org/gsea/index.jsp) and Enrichr (https://maayanlab.cloud/Enrichr) (77) tools.

### *In silico* analysis of the RNA-protein interactions

The protein structures of the four targets RPS3A, RPS7, RPL5, RPL26 are not reported, therefore, their sequences were retrieved from the “uniport” by using their id numbers P61247, P62081, P46777, and P61254, respectively. Furthermore, the i-tasser module was used to build their 3D structures. Based on their c-score, the best model was selected and evaluated through Ramachandran plots. Furthermore, the energy minimization of each protein was carried out using AMBER tool. The 3dRNA tool is used to generate the 3D of RNA sequence. For Protein-RNA docking, the HDOCK was used with the standard cut-offs. In each docking the top complex was picked for further analysis. The energy minimized complex structures were used for their quantitative and thermodynamic analysis.

### Luciferase reporter assay, Western blot analysis, RNA isolation and quantitative real time PCR (RTq-PCR)

To perform the dual luciferase reporter assay, the HEK 293T cells were seeded at 70% confluency and next day, 25nM siRNA were transfected in respective wells using 0.35µl of Dharmafect transfection reagent as per manufacturer’s instructions (GE Healthcare Dharmacon Inc. Co, USA). 12 hours post transfection of siRNA, media was changed followed by transfection of the dual luciferase reporter plasmids (pRLFF/luc87, 1μg/well) using lipofectamine 2000 at 1:1 (wt/v) ratio. 8 hours later, media was changed and cells were incubated for 48 hours. Cells were washed with ice cold PBS and used for luciferase assay using the dual glow Luciferase assay kit, following manufacturer’s protocol (Promega, Wisconsin, USA). The Firefly-Luc values were divided by that of the Renilla-Luc or divided by the percentage cell-viability, as specified in the results and graphs were plotted in GraphPad prism or Microsoft Excel. Values are mean (± SEM) of three independent experiments done in triplicates. Western blot was done as described (32). Total cellular RNA was isolated using the TRI reagent (MRC Inc., Cincinnati, USA) as per manufacturer’s protocol. cDNA was synthesized using random hexamers and a commercial cDNA synthesis kit, following manufacturer’s protocol (Firescript cDNA synthesis kit, Solis BioDyne, Tartu, Estonia). RTq-PCR was done by SYBR green detection method using a commercially available kit (5X Hot Firepol Evagreen qPCR mix plus, Solis BioDyne, Tartu, Estonia). Following primers were used to detect the levels of g1-HEV RNA, HEV IRESl RNA, Renilla-Luc RNA and control RNA (618-738 nt in g1-HEV genome): g1-HEV FP: (5’ CGGCCCAGTCTATGTCTCTG 3’), g1-HEV RP: (TAGTTCCTGCCTCCCAAAAG 3’); HEV IRESl FP (5’ TATACTCGAGGGTGCCGATCGGTCCC 3’), HEV IRESl RP (5’ TATACCATGGCATCTGGCAGCAAGCTCAG 3’); R-Luc FP (5’ CTTCTTATTTATGGCGACATGTT 3’) and R-Luc RP (5’ GCCTGATTTGCCCATACC AATA 3’); Control RNA FP (5’ GCCCCCTGGCACATA 3’) and Control RNA RP (5’ GAGCGCAGGTTGGAAA 3’).

### Ribosome fractionation assay

Ribosome fractionation was done as described (43, 44). HEK 293T cells were seeded at 70% confluency in 100 mm dishes and transfected with the pGL3-control RNA, pSuper-HEV IRESl and pRL-TK plasmid DNA (12µg/dish) using lipofectamine 2000 (1:1 ratio). 48 hours post-transfection, 100 µg/ml of cycloheximide (CHX) or 0.5µg/ml of puromycin (Puro) was added to the culture medium and incubation was continued for 30 minutes at 37^0^C, with 5% CO_2_. Next, cells were washed three times with ice cold PBS (containing 100 µg/ml CHX, in case of CHX treatment) followed by trypsinization and centrifugation for 5 minutes at 500×g. Supernatant was discarded and the cell pellet was resuspended in 1ml polysome extraction buffer (PEB, 20mM Tris-HCl, pH 7.5, 100mM KCl, 5mM MgCl_2_, 0.5% Nonidet P-40, 100µg/ml CHX, 1X protease inhibitors and 1:1000 dilution of RiboLock RNase inhibitor), incubated on ice for 30 minutes with occasional mixing, centrifuged at 12000×g for 20 minutes at 4^0^C. Total protein and RNA concentration in the clarified lysate was measured by Bradford assay or Nanodrop spectrophotometry, respectively. Aliquots of the lysate were used for cDNA synthesis, western blot analysis and ribosome fractionation.

A linear 10% - 50% sucrose density gradient was prepared for ribosome fractionation, one day before the polysome fractionation experiment. Stock solutions of 10%, 20%, 30%, 40% and 50% sucrose were prepared using 2.2M sucrose and 10x salt solution (1000 mM NaCl, 200mM Tris-HCl, pH 7.5, 50 mM MgCl_2_) in Diethyl Pyrocarbonate (DEPC)-treated water. 2.2 ml of 10% stock solution of sucrose was added to the bottom of a thin-wall 12.5 ml ultracentrifuge tube, followed by addition of 2.2 ml each of 20%, 30%, 40%, and 50% sucrose solutions. Tubes were stored at 4^0^C, overnight for linearization. Next day, equal amount of clarified lysate was loaded to the top of the tubes and weights were equalized. Tubes were run in a SW40Ti swinging bucket rotor at 190000×g for 90 minutes at 4^0^C, using a Beckman ultracentrifuge (Beckman Coulter, California, USA). After centrifugation, tubes were set for fractionation in the automated fraction collection unit and 24 fractions of 500 µl each were collected. Profile of RNA in the fractions was captured as an indicator of different ribosome fractions (shown in Fig. S4A). Alternate fractions (1–23) were selected for further analysis of RNA and protein distribution profiles.

### M^7^G capped-RNA and biotinylated-RNA pull down assay

For m^7^G capped RNA pulldown, HEK 293T cells were seeded in 60 mm dishes at a confluency of 70%. Next day, cells were cotransfected with 2µg each of the pSuper-HEV IRESl and pRL-TK plasmids and cells were incubated for 48 hours in DMEM media supplemented with 10% FBS in a 37^0^C incubator supplied with 5% CO_2_. Next, cells were lysed in 1ml TRI reagent and RNA was extracted as per manufacturer’s instructions (MRC Inc, Cincinnati, USA). 20µg of total RNA was added to the binding buffer [1% Triton X-100, 150mM NaCl, 2mM EDTA (pH 8.0), 20mM Tris-HCl (pH 8.0)], 20 µl monoclonal anti-m^7^G cap antibody (Merck, Massachusetts, USA) and 1µl RNAse inhibitor (RNAsin, Promega, Wisconsin, USA) was added to that and samples were incubated overnight at 4°C with continuous mixing. 25 µl Protein G-PLUS-Agarose beads (Santa Cruz Biotechnology, Santacruz, USA) were washed twice in the binding buffer and added to the samples, followed by one hour incubation. Samples were centrifuged at 1000 rpm for 1 minute at 4^0^C. Supernatant containing unbound RNA was collected, RNA was extracted using TRI reagent and stored at −80^0^C. Beads were washed four times in ice cold low salt wash buffer [1% SDS, 1% Triton X-100, 150mM NaCl, 2mM EDTA (pH 8.0), 20mM Tris-HCl (pH 8.0)] and once in high salt wash buffer [1% SDS, 1% Triton X-100, 500mM NaCl, 2mM EDTA (pH 8.0) 20mM Tris-HCl (pH 8.0)] and twice with Tris-EDTA buffer (pH 8.0). Finally, beads were re-suspended in the elution buffer (1% SDS, 0.1M NaHCO3), incubated for 5 minutes at 4^0^C on a rocker, followed by centrifugation at 1000 rpm for 1 minute at 4^0^C and collection of the supernatant. RNA was isolated from the supernatant using TRI reagent and all samples were simultaneously processed for RT-qPCR.

For biotinylated RNA pulldown assay, 10^7^ Huh7 cells were lysed in the binding buffer [1% Triton X-100, 0.5% NP-40, 150mM NaCl, 2mM EDTA (pH 8.0), 20mM Tris-HCl (pH 8.0), 1X complete protease inhibitor cocktail, 40 units RNAse inhibitor] and clarified lysate were incubated with biotinylated control RNA [control RNA (B^+^)], non-biotinylated HEV IRESl RNA [HEV IRESl RNA (B^-^)], biotinylated HEV IRESl RNA [HEV IRESl RNA (B^+^)] or mock lysate (Cell lysate without RNA) for 1 hour on a flip-flop rocker at 4^0^C. Streptavidin magnetic beads were washed twice in binding buffer and added to the RNA-protein mixture, followed by incubation for 1 hour, under similar condition. Streptavidin bound RNA-protein complex were washed two times in ice cold low salt wash buffer [1% SDS, 1% Triton X-100, 150mM NaCl, 2mM EDTA (pH 8.0), 20mM Tris-HCl (pH 8.0)] and once in high salt wash buffer [1% SDS, 1% Triton X-100, 250mM NaCl, 2mM EDTA (pH 8.0) 20mM Tris-HCl (pH 8.0)] and once with Tris-EDTA buffer (pH 8.0). 50% of the beads were re-suspended in the elution buffer (1% SDS, 0.1M NaHCO_3_), incubated for 5 minutes at 4^0^C on a rocker, followed by collection of the supernatant for checking the presence of RNA by formaldehyde-agarose gel electrophoresis. Rest of the beads were mixed with 2X Laemmli buffer, incubated at 95^0^C for 5 minutes, followed by collection of the supernatant for western blot analysis using various antibodies.

### Statistical analysis

Data are shown as mean ± standard error of mean (SEM) of three independent experiments. *P*-values were calculated by two tailed Student t-test (paired two samples for means).

## Acknowledgments

RNA motif plasmid cloning backbone (pRMB) and BASU RaPID plasmid were kindly gifted by Prof. Paul Khavari. We thank Prof. Marvin Wickens for providing the Y3H assay system. We thank Prof Charles M. Rice for providing the g3A HCV replicon. We thank Mr. Dipanka Tanu Sarmah and Dr. Samrat Chatterjee for data analysis using “Enrich R”. We thank Dr. Saumya Anang and Dr. Chandru Subramani for helping in the initial experiments. We appreciate continuous support by the technical staff of RCB mass spectrometry facility in data collection. This study was funded by the Science and Engineering Research Board (SERB), Govt of India, grant to MS and THSTI core grant to MS. SK is supported by a senior research fellowship from the Department of Biotechnology, Govt of India, RV and SS are supported by a senior research fellowship from the Council of Scientific and Industrial Research, Govt of India.

## Supplementary Figures and Tables

**Figure S1.**
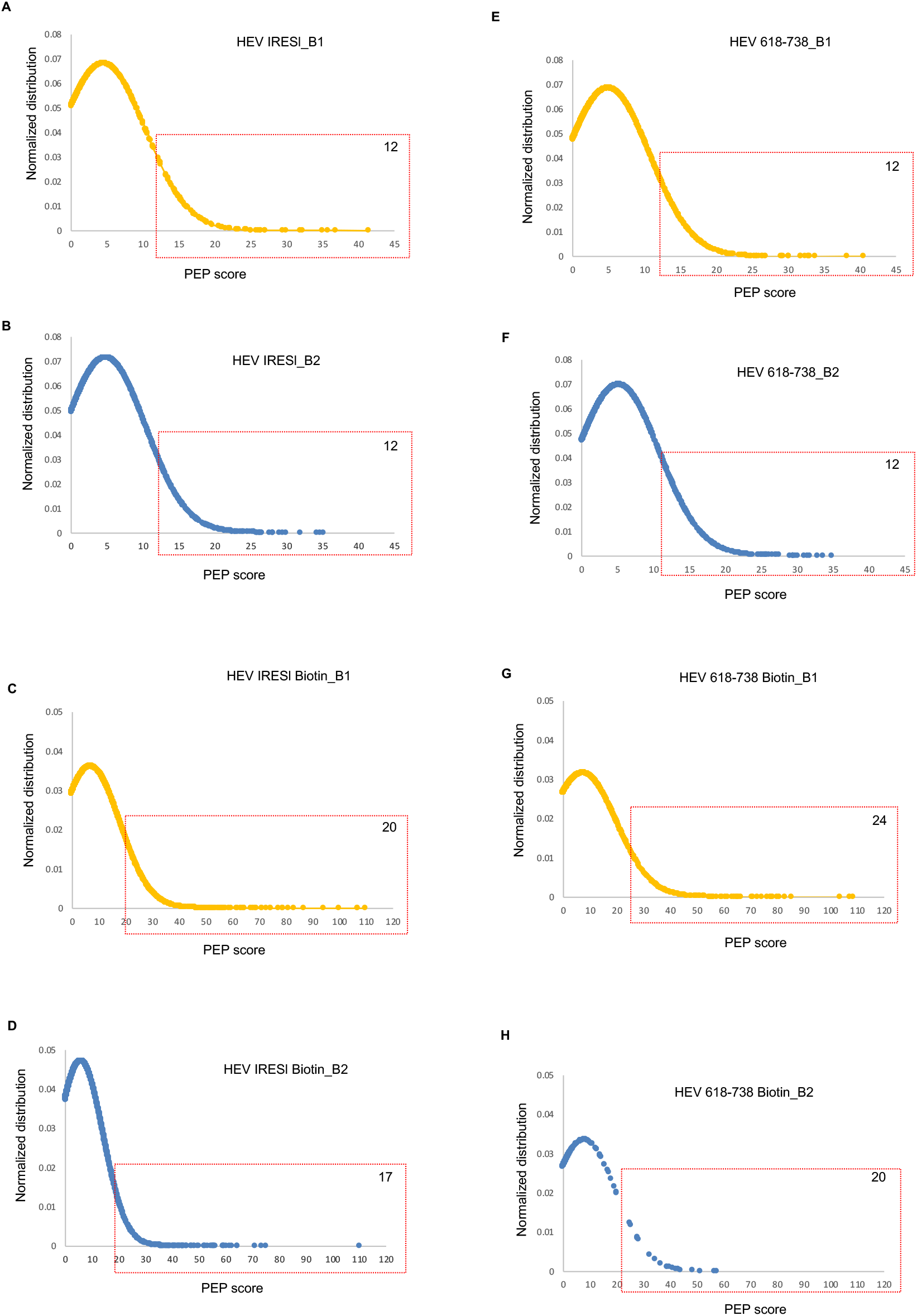
Normalized distribution plots of the peptides identified by LC-MS/MS. (A-H) Normalized distribution plots showing all peptides identified in LC-MS/MS against their respective PEP scores. B1 and B2 denote biological replicates of the same samples. PEP score range considered for protein identification is shown by a dotted-lined box.

**Figure S2.**
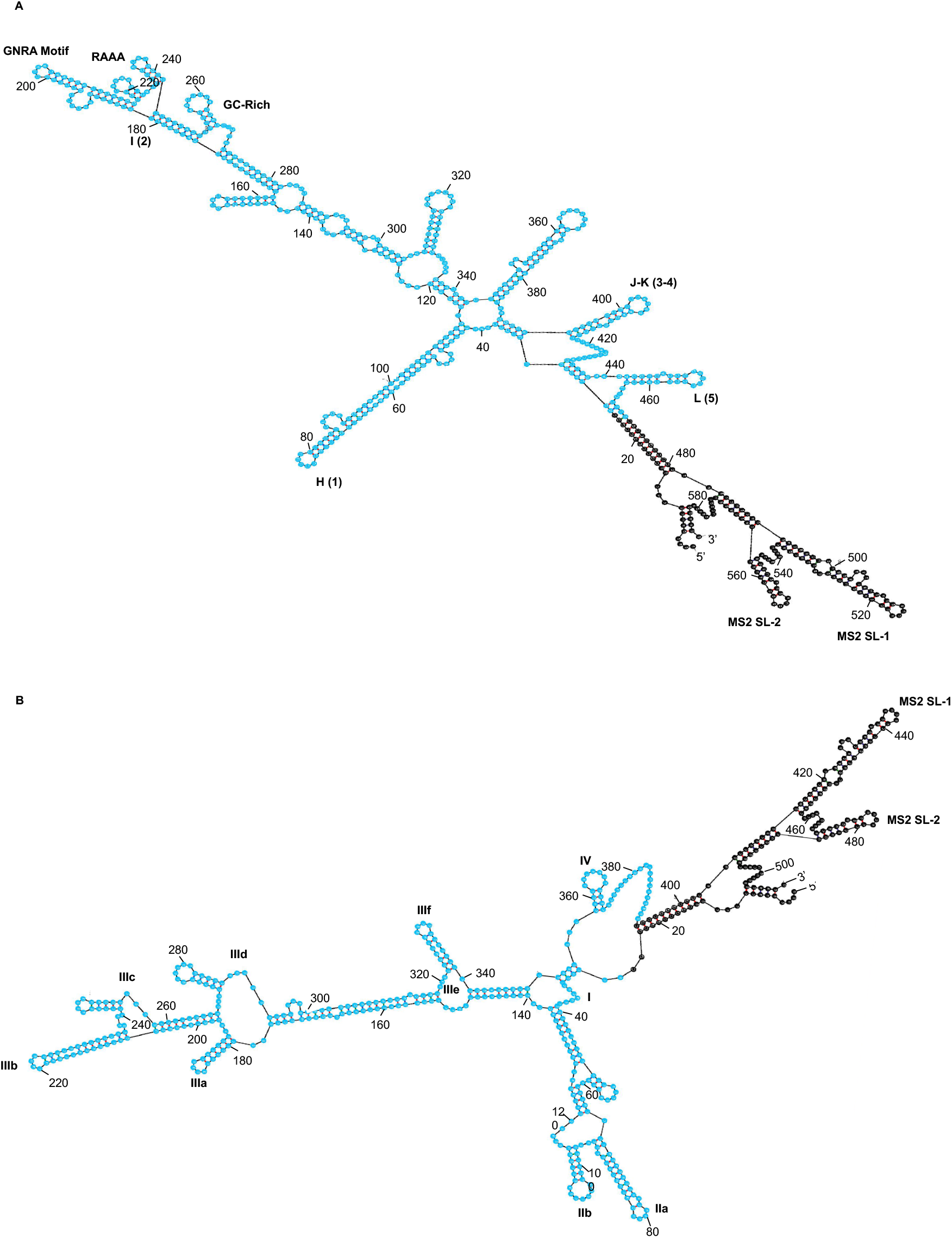
Predicted secondary structure of the FMDV and HCV IRES RNA. (A) Predicted secondary structure of the FMDV IRES RNA (shown in blue) fused to the MS2-coat protein-binding RNA. MS2 SL-1 and MS2 SL-2 represent MS2 coat proteinbinding RNA motifs. Five SLs of the IRES are numbered H-L (1-5) and span positions 26113, 114-344, 343-418 and 417-470, respectively. GNRA, RAAA, GC-rich motifs are as indicated. (B) Predicted secondary structure of the HCV IRES RNA (shown in blue) fused to the MS2-RNA. MS2 SL-1 and MS2 SL-2 represent MS2 coat protein-binding RNA motifs. Four stem-loops (SL) of the IRES are numbered as I, II, III (a-e) and IV.

**Figure S3.**
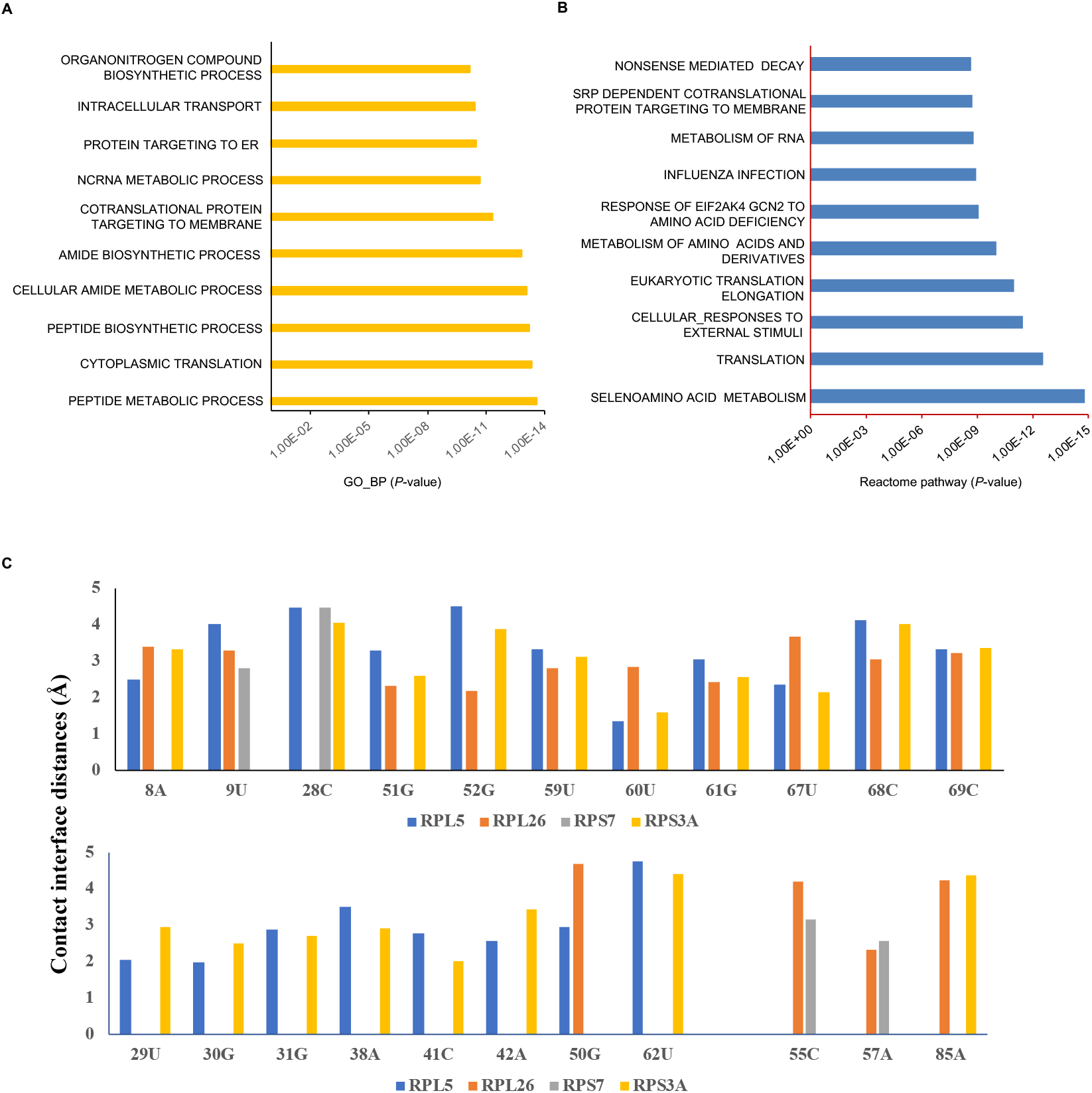
Bioinformatics analysis of the HEV IRESl RNA-host protein interaction. (A) Graphical representation of the top 10 Biological processes (sorted by P-values) enriched in the HEV IRESl RNA-protein interactome, analyzed by the GSEA tool (B) Graphical representation of the top 10 Reactome pathways (sorted by P-values) enriched in the HEV IRESl RNA-protein interactome, analyzed by the GSEA tool. (C) Quantitative pair-wise contact interface distance calculation between the HEV IRESl RNA and the indicated ribosomal proteins. The pair-wise interface residues contacts (measured in the form of distance in Å) was calculated for each complex. The common contact nucleotides of the HEV IRESl RNA are shown in the upper and lower panels, respectively.

**Figure S4.**
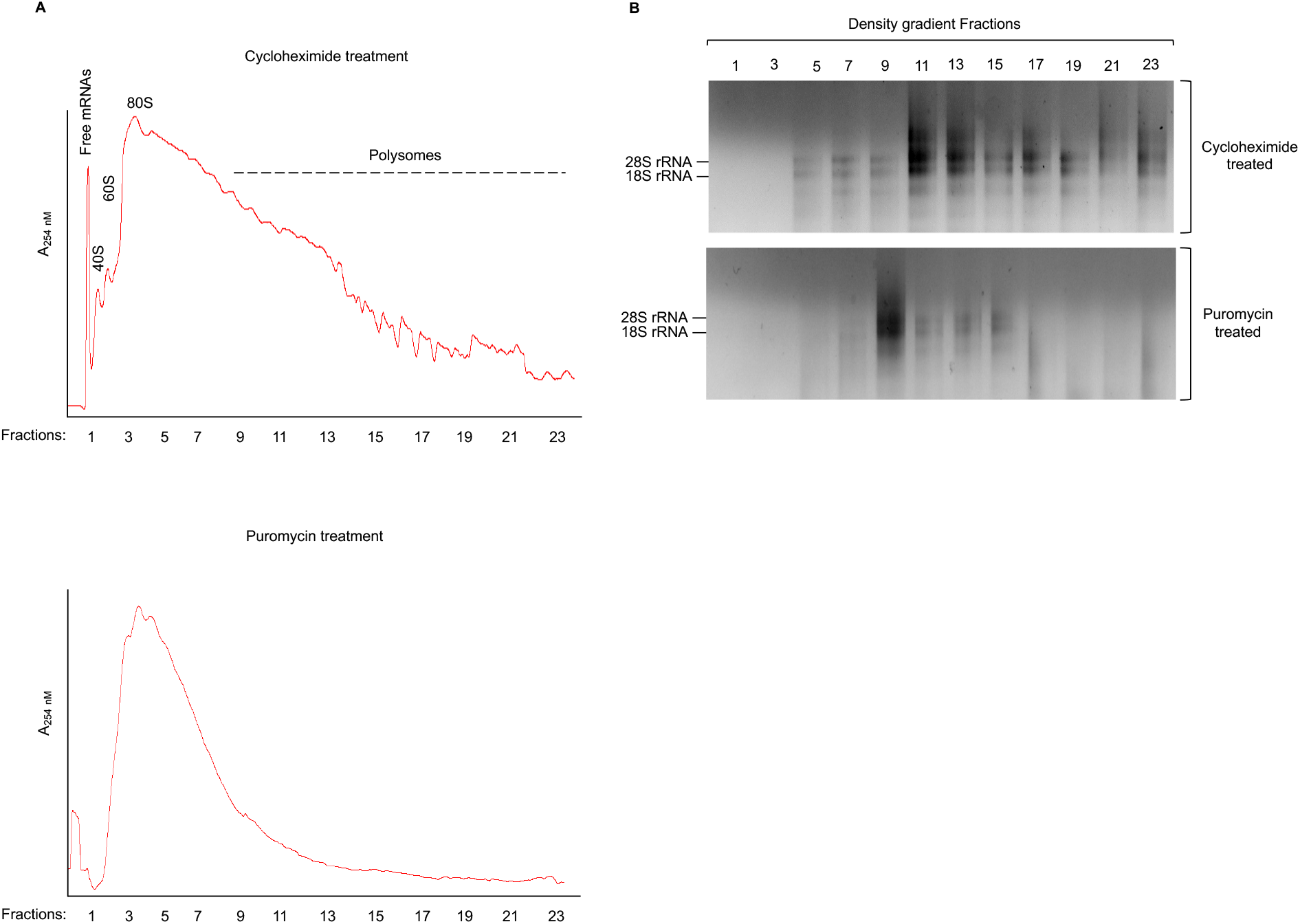
Ribosome fractionation profiles of cycloheximide or puromycin treated Huh7 cells. (A) HEK 293T cells were treated with cycloheximide for 30 min, followed by preparation of the cytosolic extract and ultracentrifugation on 10-50% sucrose gradients. Equal volume fractions were collected from the top in an automated fraction collector. Real-time profile of the RNA content in each fraction was measured at A254nm and the plot generated by the application software is shown. Peaks corresponding to free mRNAs, 40S, 60S, 80S ribosome and polysome fractions are indicated based on earlier reports (upper panel). HEK 293T cells were treated with puromycin for 30 min, followed by preparation of the cytosolic extract and ultracentrifugation on 10-50% sucrose gradients. Equal volume fractions were collected from the top in an automated fraction collector. Real-time profile of the RNA content in each fraction was measured at A254nm and the plot generated by the application software is shown (lower panel). (B) Formaldehyde-agarose gel electrophoresis of total RNA isolated from the sucrose density gradient fractions of the Huh7 cells, cotransfected with the pSuper-control RNA plasmid, pSuper-HEV IRESl plasmid and pRL-TK plasmid and treated with cycloheximide (upper panel) or puromycin (lower panel).

**Table S1.**
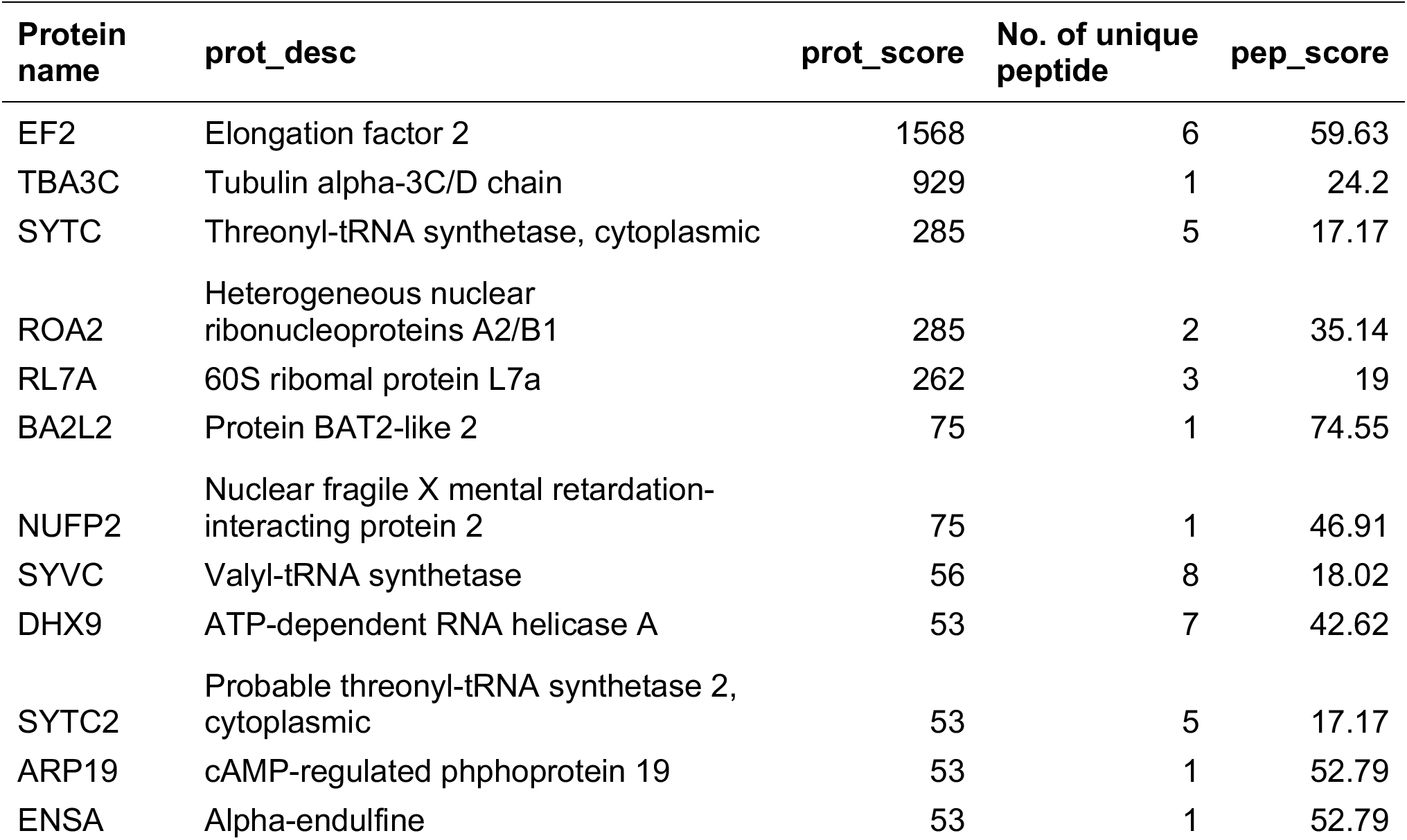
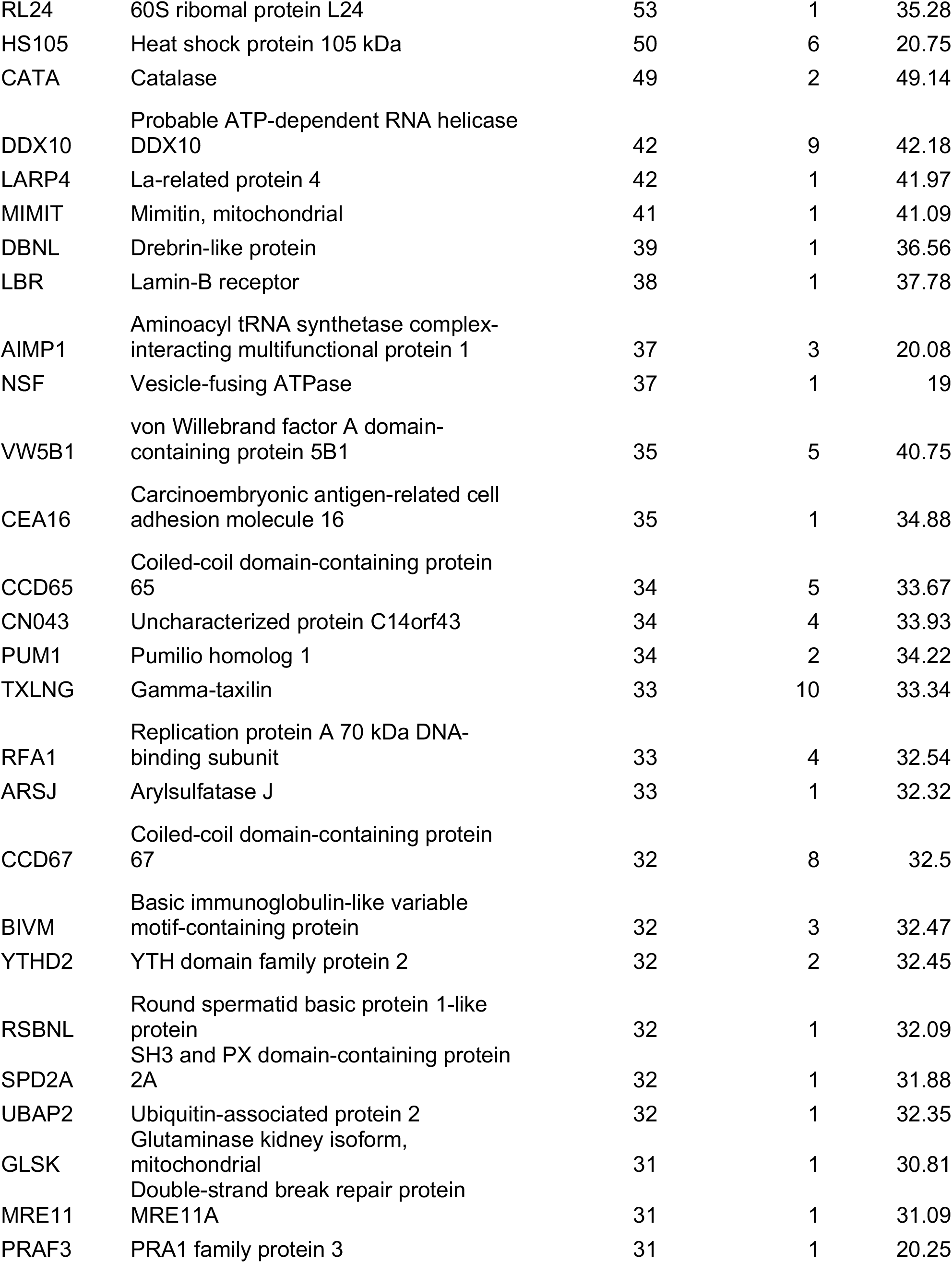
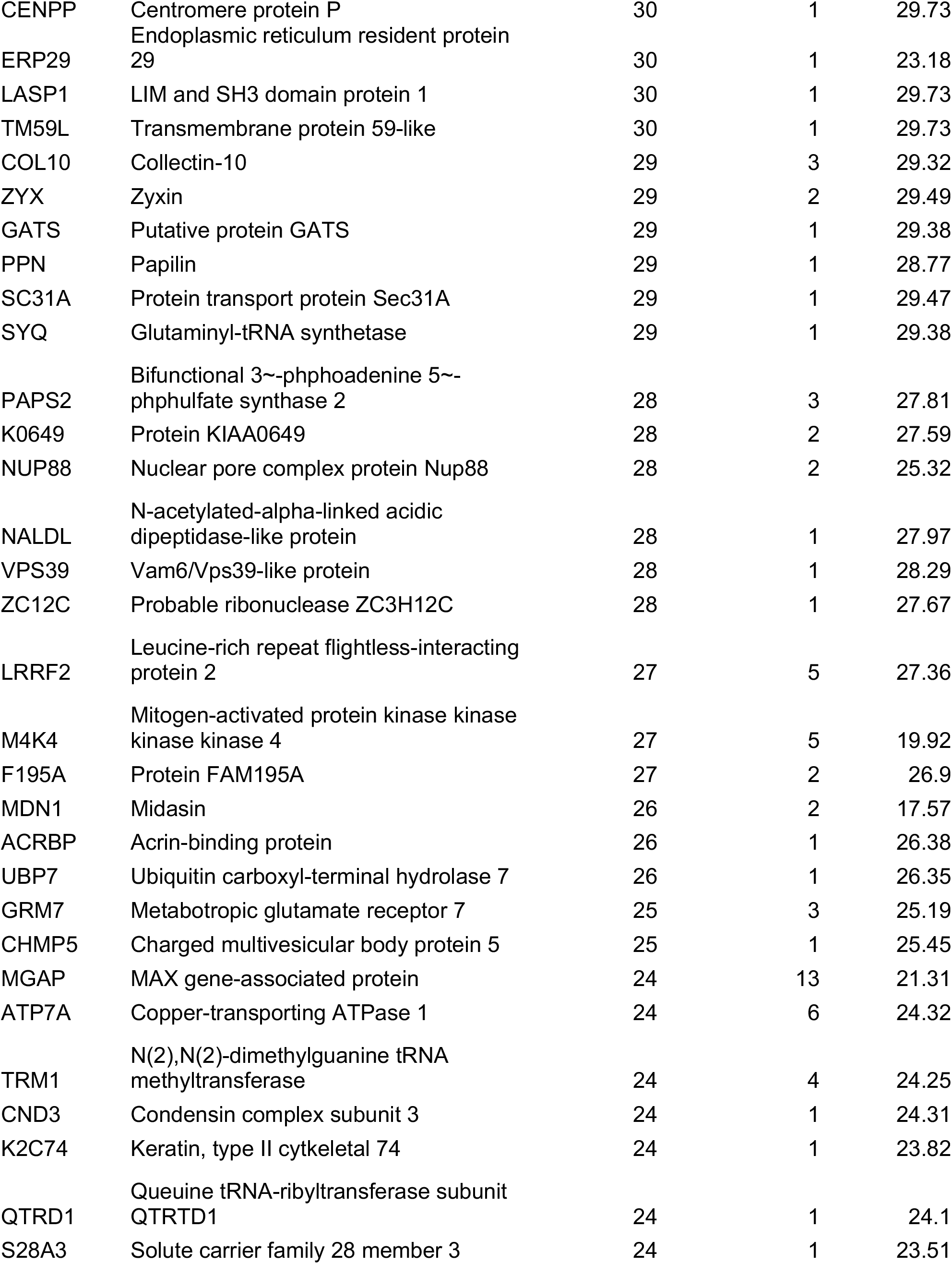
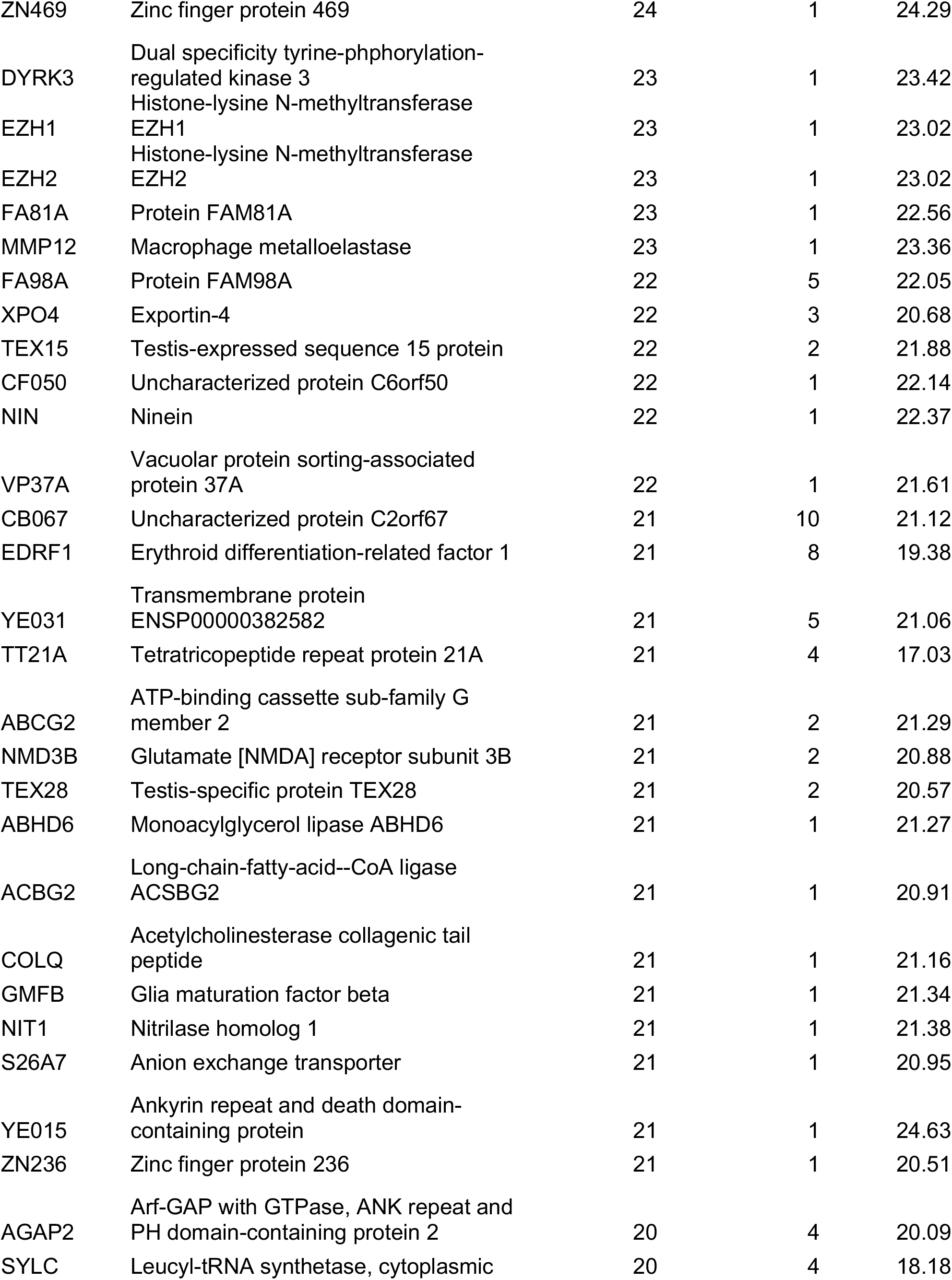
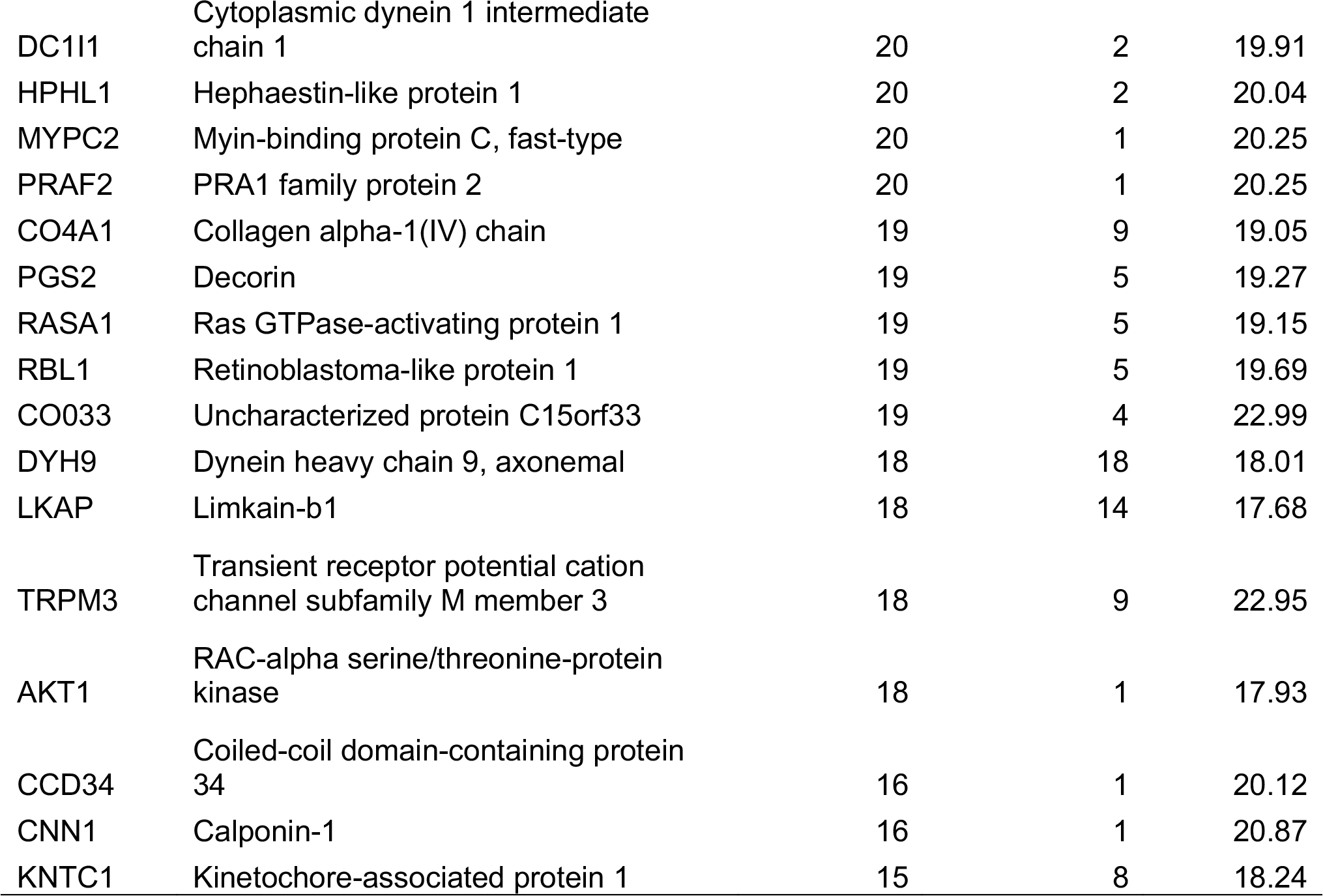
List of the HEV IRESl RNA interaction partners, identified by RaPID-LC-MS-MS.

**Table S2.**
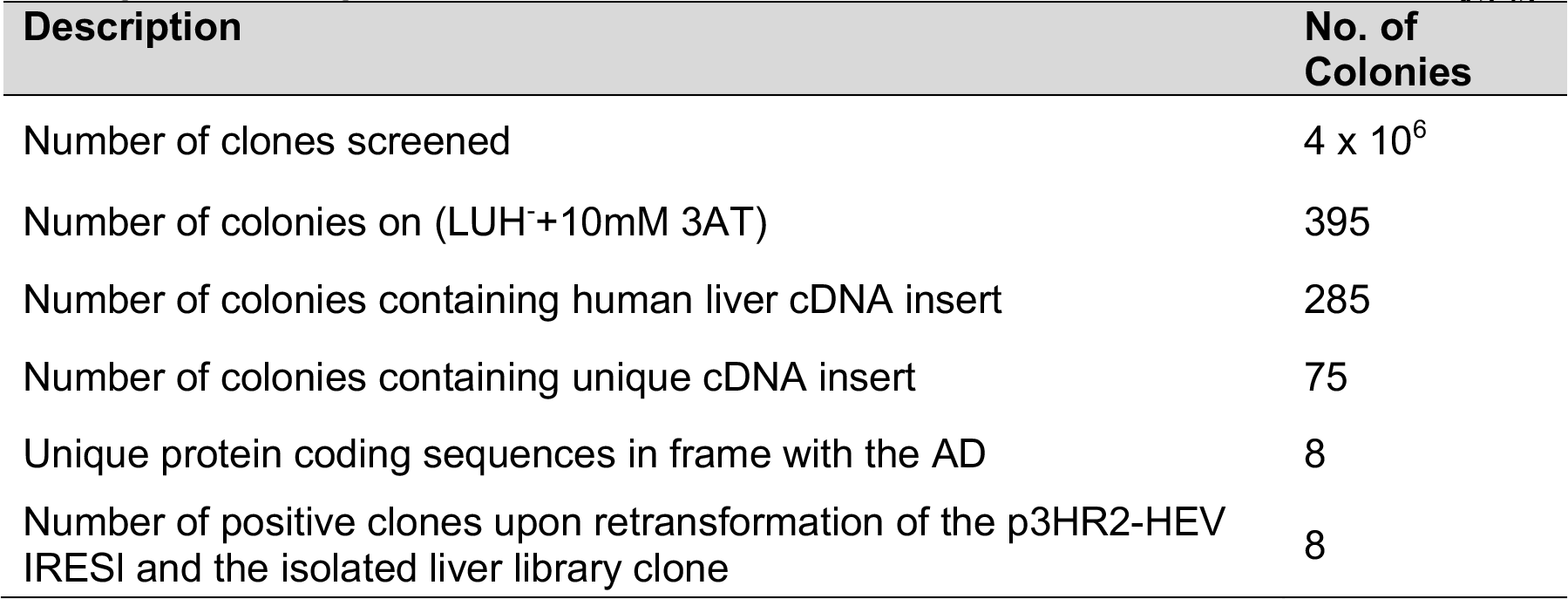
Summary of human fetal liver library screening by the Y3H^1371^ assay to identify the host factors that interact with the HEV IRESl element.

**Table S3.**
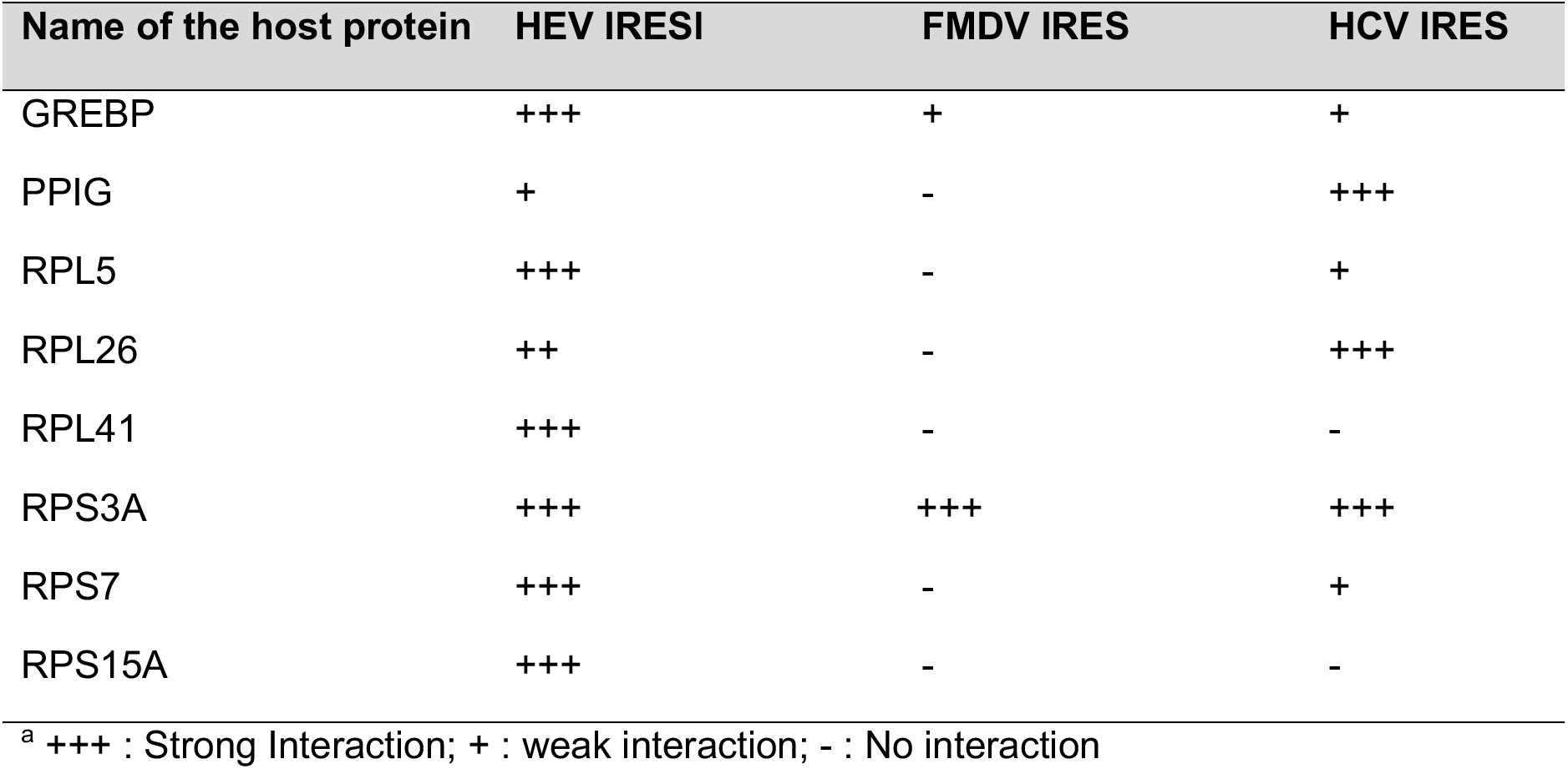
Analysis of interaction of the HEV IRESl RNA-interacting host proteins (found in Y3H library screening) with the FMDV and HCV IRES RNA^a^.

| Name of the host protein | HEV IRESI | FMDV IRES | HCV IRES |
| --- | --- | --- | --- |
| GREBP | +++ | + | + |
| PPIG | + | - | +++ |
| RPL5 | +++ | - | + |
| RPL26 | ++ | - | +++ |
| RPL41 | +++ | - | - |
| RPS3A | +++ | +++ | +++ |
| RPS7 | +++ | - | + |
| RPS15A | +++ | - | - |

<sup>a</sup> +++ : Strong Interaction; + : weak interaction; - : No interaction

